# From Sensory to Perceptual Manifolds: The Twist of Neural Geometry

**DOI:** 10.1101/2023.10.02.559721

**Authors:** Heng Ma, Longsheng Jiang, Tao Liu, Jia Liu

**Affiliations:** Department of Psychological and Cognitive Sciences & Tsinghua Laboratory of Brain and Intelligence, Tsinghua University, Beijing 100084, China; Beijing Academy of Artificial Intelligence (BAAI), Beijing 100085, China

**Author notes:** These authors contributed equally: Heng Ma, Longsheng Jiang, Tao Liu.

**Keywords:** motion-induced illusory contour, linearly inseparable problem, neural geometry, sensory manifold, perceptual manifold, nonlinear mixed selectivity

## Abstract

Classification constitutes a core cognitive challenge for both biological and artificial intelligence systems, with many tasks potentially reducible to classification problems. Here we investigated how the brain categorizes stimuli that are not linearly separable in the physical world by analyzing the geometry of neural manifolds in high-dimensional neural space, formed by macaques’ V2 neurons during a classification task on the orientations of motion-induced illusory contours. We identified two related but distinct neural manifolds in this high-dimensional neural space: the sensory and perceptual manifolds. The sensory manifold was embedded in a 3-D subspace defined by three stimulus features, where contour orientations remained linearly inseparable. However, through a series of geometric transformations equivalent to twist operations, this 3-D sensory manifold evolved into a 7-D perceptual manifold with four additional axes, enabling the linear separability of contour orientations. Both formal proof and computational modeling revealed that this dimension expansion was facilitated by nonlinear mixed selectivity neurons exhibiting heterogeneous response profiles. These findings provide insights into the mechanisms by which biological neural networks increase the dimensionality of representational spaces, illustrating how perception arises from sensation through the lens of neural geometry.

## Introduction

Imagine a person trying to identify various objects such as a cup, a book, and a pen on a clustered desk. The brain must process complex and overlapping sensory inputs with variations in light, angles, and occlusions by neurons encoding different features of the objects and their combinations to ultimately classify each object accurately (Biederman, 1987; DiCarlo et al., 2012; Chung & Sompolinsky, 2018). One primary challenge in this classification task lies in the prevalence of linearly inseparable problems, where perfectly segregating data points into their respective classes using a linear boundary is infeasible (Minsky & Papert, 1969; Fusi et al., 2016). Machine learning algorithms often resorts to complex, nonlinear decision boundaries to address this issue, employing techniques like kernel methods and deep learning (Schölkopf & Smola, 2002; Williams & Rasmussen, 2006). Here we asked how the brain addresses linearly inseparable problems present in the physical world from the perspective of neural geometry (Seung & Lee, 2000; DiCarlo & Cox, 2007; Chung & Abbott, 2021) constituted by the collective activity of large groups of neurons, an approach recently applied to various domains such as vision (Froudarakis et al., 2020; Dyballa et al., 2023), memory (Xie et al., 2022; She et al., 2024), decision (Okazawa et al., 2021; Flesch et al., 2022; Genkin et al., 2023), navigation (Gardner et al., 2022; Low et al., 2023) and motor execution (Churchland et al., 2012; Gallego et al., 2017; Gallego et al., 2018; Russo et al., 2020) to explore their characteristics in a high-dimensional neural space.

To do this, we first designed a set of stimuli that cannot be linearly classified along one stimulus feature. Specifically, we employed a visual illusion of motion-induced contours (MIC, Marcar et al., 2000; Mysore et al., 2006; Chen et al., 2014), where moving dots generate an illusory contour, either left- or right-tilted, at the center of a circular viewing area (Fig. 1B).

**Figure 1.**
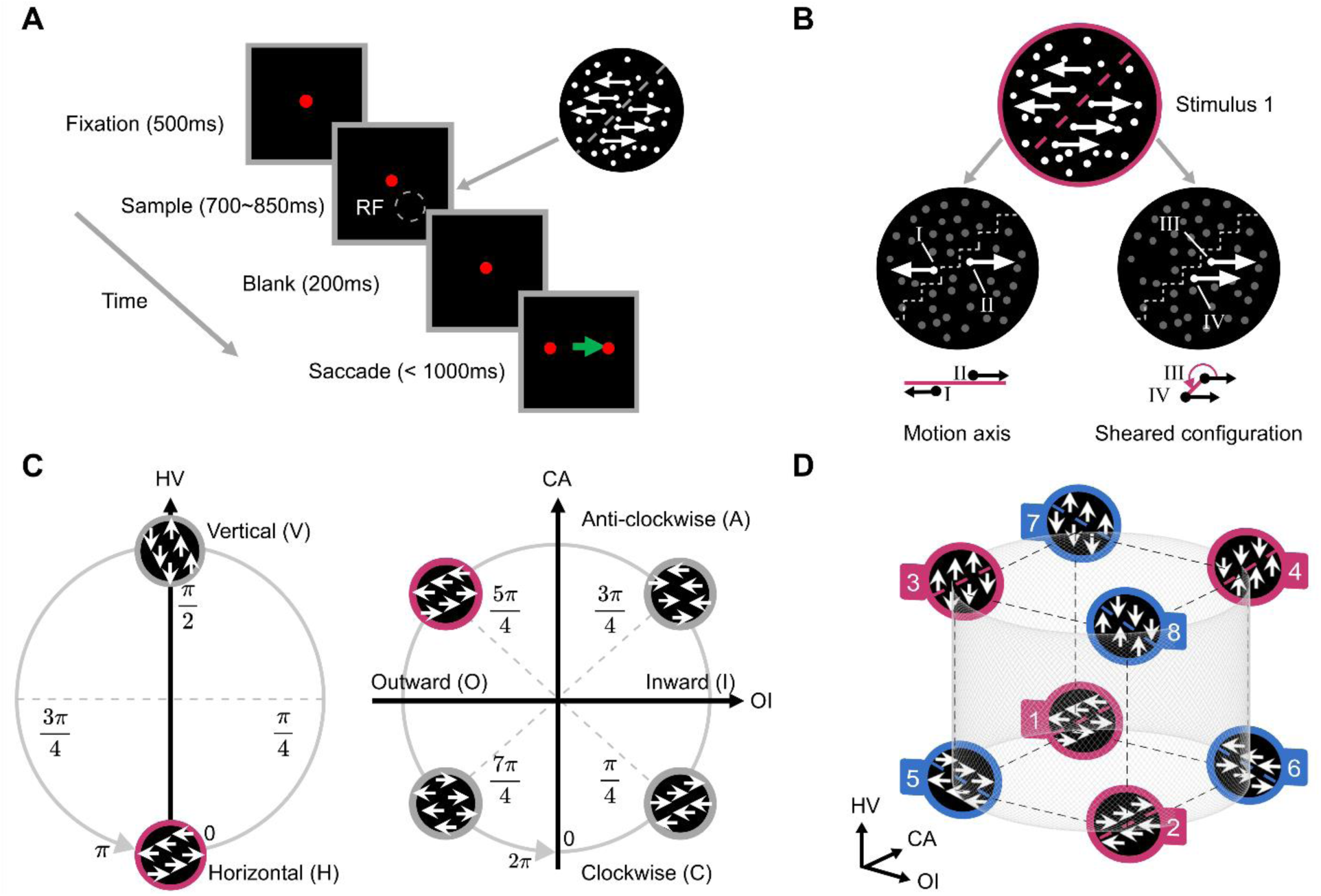
Stimulus Space. (A) Left: Discrimination task on contour orientations. Monkeys were trained to make saccades according to contour orientations. Right: A right-titled contour orientation formed by surrounding moving dots. This figure is modified from Ma et al, 2021. RF: Array population receptive field. (B) Stimulus features. An MIC stimulus is defined by the motion axis of moving dots (left: dot I and II move horizontally) and the sheared configuration of the relative positioning of adjacent dots within the same half (right: dot III and IV is 5𝜋/4 in sheared configuration, which is outward and anti-clockwise-like). (C) Feature axes of stimulus space. Left: HV axis. Two values sampled from the motion directions of [0, π], representing horizontal (0) versus vertical (π/2) directions, thus forming the HV (Horizontal vs. Vertical) axis. Right: OI and CA axes. Four values sampled from the sheared configuration of [0, 2π]: π/4, 3π/4, 5π/4 and 7π/4, creating two independent axes of OI (outward versus inward) and CA (clockwise-like versus anti-clockwise-like). Note that the feature “clockwise” (and anti-clockwise) reflects the relation positions of moving dots, not the actual rotation of moving dots. Within these three feature axes, the stimulus (red) is horizontal, outward, and anti-clockwise-like. (D) Stimulus space. The eight stimuli used in the study, each labeled numerically and color-coded based on their contour orientations (red: right-tilted; blue: left-tilted). In this stimulus space, the classification of contour orientations poses a linearly inseparable problem.

There are three independent stimulus features of moving dots that determine a particular instance of MIC stimuli (Chen et al., 2014; Lu et al., 2010; Ma et al., 2021): dots move (1) either horizontally or vertically, (2) outwardly or inwardly, and (3) clockwise-like or anti-clockwise-like (Fig. 1C). Accordingly, a 3-D stimulus space is thus constructed (Fig. 1D), with each axis corresponding to a stimulus feature of moving dots. Interesting, although each stimulus feature of moving dots is linearly separable, the orientations of illusory boundary, constructed by combining these three features, become linearly inseparable in this stimulus space. That is, MIC stimuli sharing the same contour orientation (e.g., right-titled, red) are interspersed among those with the opposing orientation (i.e., left-tilted, blue). Therefore, the classification of contour orientations presents a linearly inseparable problem in the physical world.

In the mental world, however, the linear classification of contour orientations becomes feasible, as previous neurophysiological studies have revealed that single neurons in macaques’ V2 exhibit selectivity for cue-invariant contour orientations (Marcar et al., 2000; Ma et al., 2021). Accordingly, we analyzed neuronal activity data recorded in the V2 area (Ma et al., 2021), aiming to elucidate how the linearly inseparable problem in the stimulus space becomes linearly separable in the high-dimensional neural space constructed by the collective activity of V2 neurons. Importantly, because the contours do not exist in the physical world, being solely created by the brain when integrating individual motions and interpreting symmetrical movement patterns, we specifically examined two types of neural manifolds embedded in this high-dimensional space: the sensory and perceptual manifolds. The sensory manifold, which arises from the sensation process, directly responds to physical stimuli without involving interpretation and thus provides the raw sensory data that the brain uses to build perceptual experiences. Given its correspondence to external stimuli, contour orientations likely remain linearly inseparable in the sensory manifold. In contrast, the perceptual manifold reflects the brain’s effort to interpret and make sense of the sensory data, where neural states for MIC stimuli may be separately clustered based on contour orientations. Therefore, understanding the geometric difference between the sensory and perceptual manifolds and further deriving potential mechanisms that create such a difference provides a new perspective on how perception arises from sensation.

## Results

### Stimulus space and linearly inseparable problems

Two macaque monkeys were trained to perform a classification task by executing a saccade towards the side corresponding to the orientation of an MIC (Fig. 1A). The MIC stimuli used in this study were generated by moving dots, with their movement determined by two parameters. As illustrated in in Fig. 1B, dots in opposing halves moved in opposite directions along a motion axis (e.g., dot I and dot II in Fig. 1B, left). The motion axis could rotate within a range of [0, 𝜋], creating a circular structure (Supplementary Fig. 1A). For dots within the same half, one dot (e.g., dot III) either led or trailed another (e.g., dot IV) (Fig. 1B, right), quantified by the angular location of one dot relative to another, namely sheared configuration (Supplementary Fig. 2A & 2B). The sheared configuration varies periodically over [0, 2𝜋] (Supplementary Fig. 1B). Accordingly, these two periodic parameters together define a stimulus manifold with a torus topology (Supplementary Fig. 3, left), where each point represents a stimulus generated by a unique combination of the motion axis and sheared configuration. In this study, we only used a subset of this stimulus manifold. Specifically, we first projected the 2-D circular structure of the motion axis onto the 1-D axis, selecting values of 0 and 𝜋/2 (Fig. 1C, left), while retaining the 2-D circular structure of the sheared configuration (Fig. 1C, right). Consequently, the subset manifold has a cylindrical topology (Supplementary Fig. 3, right). Within this cylindrical manifold, we sampled two values, 0 and 𝜋/2, referred to as the HV (horizontal versus vertical) and four values from the sheared configuration (𝜋/4, 3𝜋/4, 5𝜋/4 and 7𝜋/4, Fig. 1C, right), defining two orthogonal axes: OI (outward versus inward) and CA (clockwise-like versus anti-clockwise-like). These three axes of HV, OI, and CA were used to construct a 3-D Euclidean space embedded in the stimulus manifold (Fig. 1D), referred to as the stimulus space. Eight stimuli used in this study were positioned at the vertices of an inner cube tangent to the cylindrical manifold.

In this stimulus space, MIC stimuli with one contour orientation formed by the combination of different values in these three axes (e.g., right-titled, No.1 – No.4, red) were interspersed with stimuli of the opposite orientation (i.e., left-tilted, No. 5 – No.8, blue) (Fig. 1D; see Supplementary Movie. 1 for the 8 stimuli). This interspersion resulted in contour orientations being linearly inseparable within this stimulus space, presenting a significant challenge for classification. However, both macaques adeptly performed the task with an accuracy exceeding 90% at 100% motion coherence (Ma et al., 2021). Next, we investigated how neurons in macaques’ V2 tackle this linearly inseparable problem particularly from the perspective of neural geometry.

### Sensory and perceptual manifolds

To investigate how the brain addressed this linearly inseparable problem, we utilized neural activity data from V2 neurons of two monkeys (93 neurons in total) to construct a high-dimension neural space (see Methods; for neurons’ receptive fields and selectivity, see Supplementary Fig. 4). First we utilized linear support vector machines (SVM) to analyze the collective activity to all the 8 stimuli to determine whether the three feature axes of the stimulus space (Fig. 1D), HV ({1,2,5,6} versus {3,4,7,8}), OI ({1,3,5,7} versus {2,4,6,8}), and CA ({2,3,5,8} versus {1,4,6,7}), could be decoded. We found that classifications along all three axes were linearly separable, with accuracies above 75% and significantly higher than the shuffled baseline (bootstrap t-tests, all 𝑝s < 0.001) (Fig. 2A). This finding was replicated with other methods such as the analysis on population average responses and the principal component analysis (Supplementary Fig. 5). Critically, the optimal separation direction vectors (determined by SVM, see Methods) for HV, OI, and CA, were mutually orthogonal. That is, the distribution of the subtended angles between these vectors was significantly more concentrated around 90° than the distribution of angles between two random direction vectors, and also significantly larger than angle distributions between the same classification vectors (Supplementary Fig. 6).

**Figure 2.**
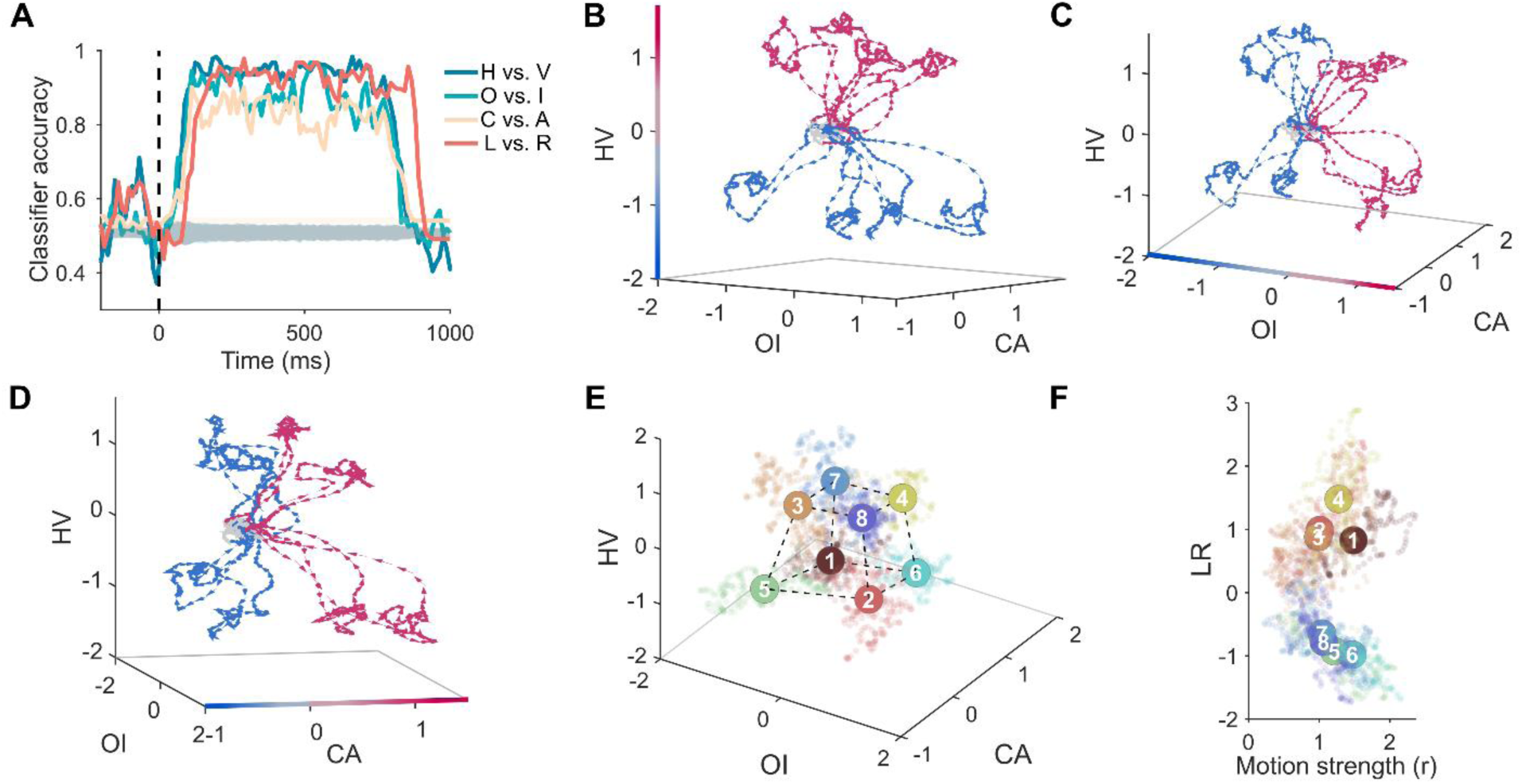
Neural geometry of sensory and perceptual manifolds. (A) Accuracies over time by linear SVM classifiers. The SVMs were trained on neural data within the time interval [-200ms, 1000ms] relative to stimulus onset. Shaded regions denote the mean SVM accuracy (±3 standard deviations) of the baselines by shuffling stimulus labels 1000 times. Individual monkey’s results are shown in Supplementary Fig. 9 E & F. (B-D) Dynamic trajectories of neural states. Changes in neural states from -200ms to 1000ms were projected onto a neural subspace defined by the HV, OI, and CA axes. Identical trajectories are presented from various perspectives to optimally illustrate the separation by these three stimulus feature axes. Blue and red colors indicate categorizations along the HV, OI, and CA axes for (B), (C), and (D), respectively. Arrows on the trajectories correspond to the progression of the neural states over time. The neural states prior to stimulus onset are indicated in gray. The motion coherence level is 7. Individual monkey’s results are shown in Supplementary Fig. 10. The time courses of neural states (-200–1000ms) projected to each axis are shown in Supplementary Fig. 11. (E) Sensory manifold. The centers of the point clouds for eight stimuli in the steady phase (200ms–500ms after stimuli onset) are connected with dashed lines. Cool and warm colors indicate the two contour orientations, respectively. All motion coherence levels were used. Individual monkey’s results are shown in Supplementary Fig. 9 A & B. (F) Dynamic trajectories of neural states along the LR axis. The point clouds follow the same color code as in panel E, with their centers marked with solid circles. All motion coherence levels were used. Note that cool and warm colors are separated along the LR axis. Motion strength (𝑟), which is the Euclidean distance between the neural states and the origin in panel E, is included for display purposes only.

To visualize the dynamic encoding of the stimuli by the neuronal population, we projected the neural states in the high-dimensional neural space into a 3-D neural subspace formed by the direction vectors for HV, OI, and CA (see Methods). Figure 2B shows that, prior to stimulus onset, neural states for the stimuli were closely clustered and inseparable. Following the onset, these states gradually spread and became completely separable along the HV axis. They then remained in a steady phase for the duration of stimulus presentation before returning to their original locations after stimulus offset (see Supplementary Movie. 2 and Supplementary Fig. 11). Similar dynamic patterns were observed for the OI (Fig. 2C) and CA (Fig. 2D) axes as well. To further illustrate the neural geometry constructed by the neural states at the steady phase, we used the activity magnitude of neurons from 200ms to 500ms post-stimulus onset as the neural states, which were then projected to this 3-D subspace (see Methods). Figure 2E shows that the neural states were located at eight vertexes of a slightly distorted cube, corresponding to the geometric relation among stimuli in the original stimulus space (Fig. 1D). This neural manifold embedded in the high-dimensional neural space is referred to as the “sensory manifold” because it directly responded to the stimulus space without involving interpretation, as contour orientations remain linearly inseparable in this manifold. To further demonstrate that this manifold was associated with sensory processing, we examined the relationship between its geometry and the intensity of sensory signals. We found that the sensory manifold was sensitive to variations in the coherence of moving dots in the stimuli, giving rise to a series of concentric cubic sensory manifolds, with higher coherence levels eliciting larger cubic manifolds (Supplementary Fig. 8). Taken together, the sensory manifold faithfully represented the stimulus space, with HV, OI, and CA serving as the axes of the neural subspace that embedded this manifold.

Based on previous studies showing that individual neurons that are sensitive to contour orientation (Marcar et al., 2000; Ma et al., 2021), we used the SVM analysis to explore axes in this high-dimensional neural space that could linearly decode contour orientations. We identified an axis that can differentiate left-tilted orientation from the right-tilted orientation, referred to as the LR axis, with an accuracy comparable to that of decoding the stimuli’s features (i.e., HV, OI, and CA) (Fig. 2A, orange line). Figure 2F illustrates the neural states along the LR axis, where contour orientations became linearly separable (for dynamic trajectories see Supplementary Fig. 11 & Supplementary Movie 2), consistent with the finding from studies on single neurons (Marcar et al., 2000; Ma et al., 2021). Critically, this LR axis was orthogonal to the three stimulus feature axes of the sensory manifold (Supplementary Fig. 6A), suggesting that the LR axis emerged not directly from the stimuli but from the interpretation of the sensory data.

Evidence supporting this conjecture comes from the analysis on the latency of the emergence of the LR axis, which occurred about 30ms after the emergence of the axes encoding the stimuli’s features (i.e., HV, OI, and CA) (Fig. 3F). This observation aligns with previous findings on visual motion segregation (Braddick, 1993; Lamme, 1995; Marcar et al., 2000; Chen et al., 2014; Hu et al., 2018), implying that extra time may be needed to form this new axis. This new neural manifold, embedded in the high-dimensional neural space including the LR axis for representing contour orientations and the axes for stimulus features (i.e., LH, OI, and CA), is therefore referred as the perceptual manifold, where the linearly inseparable problem was addressed. Next, we explored how this LR axis was formed through geometric transformation.

**Figure 3:**
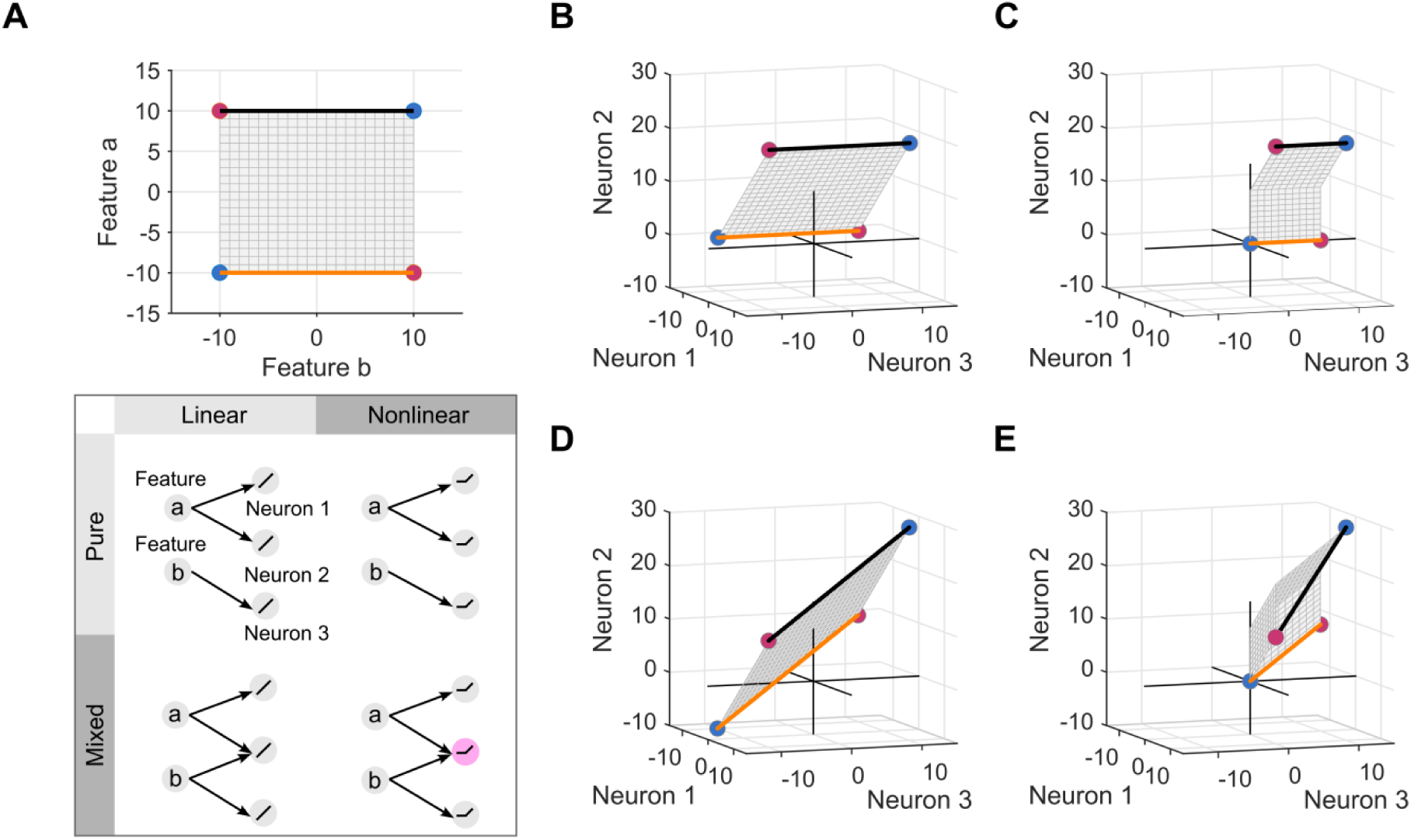
NMS neurons and twist operations. (A) Top: The 2-D feature sheet represents all possible combinations of stimulus features 𝑎 and 𝑏. Two edges of this sheet are highlighted as orange and black, and four corners are colored red and blue to indicate different categories. Bottom: Four types of neurons’ response profiles derived from the combination of input (pure versus mixed) and activation functions (linear versus nonlinear). ReLU is used for the illustration of nonlinear activation functions. The neuron marked in light purple denotes an NMS neuron. (B-E) The geometries in the neural space constructed by (B) three linear pure selectivity neurons, (C) three nonlinear pure selectivity neurons, (D) two linear pure selectivity neurons and one linear mixed selectivity neuron, or (E) two nonlinear pure selectivity neurons and one NMS neuron.

### Nonlinear mixed selectivity neurons and twist operations

Traditional solutions to linearly inseparable problems such as the XOR problem involve expanding dimensions of the original representational space (Minsky & Papert 1969; Fusi et al., 2016). Previous studies have shown that neurons with nonlinear mixed selectivity (NMS) can expand dimensions by responding to combinations of input features (Rigotti et al., 2013; Fusi et al., 2016; Kaufman et al., 2022; Tye et al., 2024) and using nonlinear activation functions (Cybenko 1989; LeCun et al., 2015) to capture higher-order interactions. In fact, all V2 neurons used in this study showed interactive responsiveness to three stimulus features (HV, OI and CA) (significant three-way interactions: minimum 𝐹(1, 2200) = 18.25, 𝑝 < 0.001), as our formal proof shows that this interactive responsiveness is a unique characteristic of NMS neurons (Supplementary Notes N.16). Therefore, these NMS neurons in the V2 area may that play a critical role in generating the LR axis through dimension expansion.

To reveal how NMS neurons expand the dimensions of representational spaces, here we examined the relationship between neurons’ response profiles and their ability to expand dimensions using a one-layer network consisting of three neurons, 𝑠_1_, 𝑠_2_ and 𝑠_3_. These neurons receive inputs consisting of two continuous features, 𝑎 and 𝑏, that form a 2-D feature sheet (Fig. 3A, top). In this context, the XOR problem is defined as categorizing points located at diagonally opposite corners (i.e., red versus blue points). The neurons in this network receive (1) either pure (either 𝑎 or 𝑏) or mixed (the combination of 𝑎 and 𝑏) inputs, and (2) possess either linear or nonlinear (e.g., ReLU) activation functions, leading to four distinct output patterns (Fig. 3A, bottom).

Neurons with pure selectivity and the linear activation function (i.e., linear pure selectivity) afford only a simple affine transformation, including rotation and shifting, of the original feature sheet (Fig. 3B). Because the rotation axis is aligned with the orange and black edges, the edges remain parallel after the transformation. For neurons with nonlinear pure selectivity, the feature sheet undergoes bending due to the nonlinear activation function (Fig. 3C). This bending occurs along an axis parallel to the orange and black edges, so the edges remain parallel after bending, and the four corner points still lie on the same plane. For neurons with linear mixed selectivity (Fig. 3D), the feature sheet experiences a complex affine transformation, with the rotation axis not aligning with the three main axes. However, even after this rotation, the edges remain parallel. In these three scenarios, neither bending nor rotation alone suffices to make the red and blue points linearly separable.

In contrast, NMS neurons perform both bending and rotation operations on the feature sheet, which make the orange and black edges no longer parallel (Fig. 3E) and thus create a new axis perpendicular to the original orange and black edges. In this newly constructed representational space, the XOR problem becomes linearly separable. The combined effects of bending and rotation operations on the 2-D feature sheet can be conceptualized as a “twist” operation. Accordingly, we refer to this unique characteristic of NMS neurons acting on the geometry of input features as the twist operation (for a similar idea, see Nogueira et al, 2023).

With this unique characteristic of NMS neurons, we further explored how the LR axis was formed from three feature axes (HV, OI, and CA) theoretically. For simplicity, we used 𝑥, 𝑦, and 𝑧 to denote the three feature axes of HV, OI, and CA, respectively. The coordinates of the points on the cylindrical manifold were denoted as (𝑥, 𝑦, 𝑧) (Fig. 4A, left). In the 𝑥*-*𝑦*-*𝑧 coordinate system, the four vertices {1,3,5,7} of the sensory manifold in the 𝑧*-*𝑥 plane exemplify a standard planar XOR problem (Fig. 4A, left). With a twist operation around the 𝑥 axis (Fig. 4A, middle), the 2-D plane is transformed into a curved surface in a 3-D space (Fig. 4A, right), creating a new coordinate axis, denoted as 𝑣 = 𝑧𝑥. Thus, the points on this curved surface have coordinates (𝑧, 𝑥, 𝑧𝑥). Further, when letting the two levels on one axis represent the true and false values of a Boolean variable, the axes are equivalent to Boolean variables (see Supplementary Methods, M.1). Accordingly, the Boolean variable 𝑉 of the new 𝑣-axis is equal to the XOR operation of the Boolean variables 𝑍 of the original 𝑧-axis and 𝑋 of the original 𝑥-axis, expressed as 𝑉 = 𝑍 ⊕ 𝑋. Therefore, the twist operation in neural geometry equals the XOR operator in logic.

**Figure 4:**
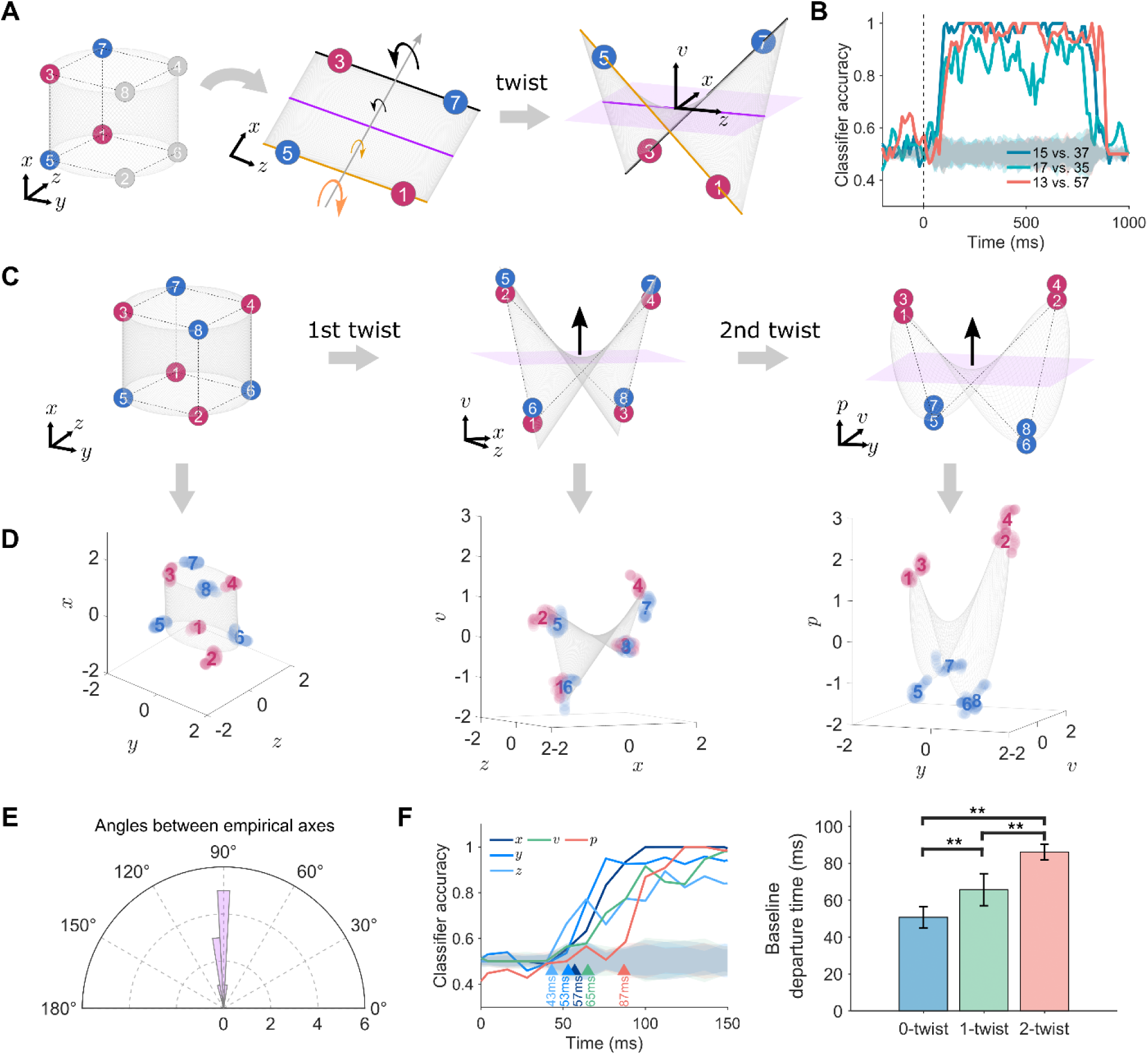
Twist operations on the classification of contour orientations. (A) Twist operations on planar XOR problems. Left: Coplanar vertices {1,3,5,7} are colored based on contour orientations, forming a standard XOR problem on the 𝑧-𝑥 plane. Middle: A twist operation around the 𝑥-axis resolves the XOR problem by rotating two parallel edges in opposite directions. Right: This twist operation generates the 𝑣 axis orthogonal to the 𝑧-𝑥 plane. (B) SVM classification accuracies over time. Shaded regions denote the mean SVM accuracy (±3 standard deviations) of the baselines by shuffling stimulus labels 1000 times. (C) Double twist operations on a cubic XOR problem. Two consecutive twists are required to correctly classify the 8 vertices according to their contour orientations. Left: Theoretical sensory manifold in the 𝑥-𝑦-𝑧 space; Middle: Intermediate manifold achieved after one twist operation on the 𝑧-𝑥 plane, with the emergence of the new 𝑣 axis. Right: Task-relevant 𝑝 axis emerges after the second twist operation on the 𝑦-𝑣 plane. (D) Neural manifolds corresponding to the theoretical manifolds achieved through twist operations. The axes of 𝑥, 𝑦, 𝑧, 𝑣 and 𝑝 in the neural space were determined by corresponding SVM classifiers. Eight point clouds of neural states corresponded to the 8 stimuli, and the centers of the clouds were used to depict the geometry of neural manifolds through linear fitting. (E) Angle distribution between 𝑥, 𝑦, 𝑧, 𝑣 and 𝑝 axes. The distribution of the subtended angles between the optimal separation direction vectors as determined by SVM was concentrated around 90°. (F) Latency of axis emergence. Left: Classification accuracies of the axes over time. The latencies of the emergence of the 𝑧, 𝑦, 𝑥, 𝑣, and 𝑝 axes (against their corresponding baselines) at 43ms, 53ms, 57ms, 65ms, and 87ms, respectively. Shaded regions denote the mean SVM accuracy (±3 standard deviations) of the baselines by shuffling stimulus labels 1000 times. Right: Latencies of the sensory axes (0-twist, blue), the intermediate axes (1-twist, green), and the task-relevant axis (2-twist, red). **: 𝑝 < 0.001. Also see Supplementary Fig. 14 C & D. For the single monkey results, see Supplementary Fig. 15.

To examine whether this theoretically derived 𝑣 axis was actually present in the high-dimensional neural space, we trained linear SVMs to classify three sets of vertex pairs among the four vertices shown in Fig. 4A. As expected, the vertex pair of {1,5} versus {3,7} and that of {1,7} versus {3,5} were linearly separable along the 𝑥 and 𝑧 axes, respectively, with accuracies greater than 75%. Critically, the vertex pair of {1,3} versus {5,7}, a standard planar XOR problem, was also linearly separable along the 𝑣 axis with accuracy greater than 75%. That is, the 𝑣 axis, theoretically derived from the twist operation, was indeed present in the high-dimensional neural space (Fig. 4B).

To examine whether this twist operation can generalize from the planar XOR problem to address the cubic XOR problem (i.e., {1, 2, 3, 4} versus {5, 6, 7, 8}, Fig. 4C, left), in addition to one twist operation equivalent to 𝑉 = 𝑍 ⊕ 𝑋 (Fig. 4C, middle), we incorporate a second twist of the manifold’s 𝑦-𝑣 projection around the 𝑦 axis to make the vertex pair of {1, 2, 3, 4} versus {5, 6, 7, 8} linearly separable (Fig. 3C, right). In this second twist, a new axis, denoted as 𝑝 = 𝑣𝑦 = 𝑧𝑥𝑦, is constructed. Logically, this axis is equivalent to 𝑃 = 𝑉 ⊕ 𝑌 = 𝑍 ⊕ 𝑋 ⊕ 𝑌, meaning the true and false values of 𝑃 is obtained by concatenating 𝑍, 𝑋, 𝑌 through two XOR operators.

To examine whether the theoretically-derived intermediate 𝑣 axis through one twist operation was indeed present in the high-dimensional neural space when considering the complete sensory manifold, we searched the 𝑧-𝑥-𝑣 subspace (Fig. 4C, middle) in the high-dimensional neural space, where the vertices {1, 3, 6, 8} was linearly separable from the vertices {2, 4, 5, 7}. The SVM analysis showed that these two sets of vertices were indeed linearly separable (Fig. 4D, middle). This finding confirms the existence of the 𝑣 axis in the high dimensional neural space. In addition, the theoretically-derived 𝑝 axis through double twist operations on the given sensory manifold is indeed the LR axis, as the 𝑝 axis was approximately parallel to the LR axis (Supplementary Fig. 13). Finally, the theoretically-derived manifolds from twist operations (Fig. 4C) closely matched the neural manifolds derived from actual neural states (Fig. 4D) (all R^2^ ≥ 0.8, for details see Methods). Note that the empirical 𝑣 and 𝑝 axes, as well as the 𝑧, 𝑥, and 𝑦 axes, were mutually orthogonal, as prescribed by twist operations (Fig. 4E). Therefore, the perceptual manifold observed in the macaques’ V2 may undergo geometric transformations equivalent to the twist operations from the sensory manifold.

Evidence supporting this conjecture comes from the analysis on the latency of the emergence of the intermediate 𝑣 axis, which should emerge after the sensory axes of 𝑧 and 𝑥, upon which the first twist operation acts, and before the task-relevant 𝑝 axis, which relies on the 𝑣 axis for the second twist. Consistent with this prediction, the empirical 𝑣 axis emerged (against the baseline) at 65ms post-stimulus onset, significantly later than the emergence of the empirical 𝑧, 𝑥, and 𝑦 axes (at 43ms, 57ms, and 53ms, respectively; bootstrap t-test, *p*s < 0.001), and yet earlier than that of the empirical 𝑝 axis (at 87ms; bootstrap t-test, *p* < 0.001) (Fig. 4F, left; for the time courses of all intermediate axes, see Supplementary Fig. 14 C & D).

Taken together, the presence of the intermediate 𝑣 axis predicted by twist operations, not by the classification task itself, suggests that the perceptual manifold is the product of mentally processing sensory data with the involvement of NMS neurons. Next, we examined the neural geometry of the perceptual manifold and its functionality.

### Dimensionality of perceptual manifold

Given the commutative nature of the equivalent logical computation, that is 𝑃 = 𝑍 ⊕ 𝑋 ⊕ 𝑌 = 𝑋 ⊕ 𝑌 ⊕ 𝑍 = 𝑌 ⊕ 𝑍 ⊕ 𝑋, we predicted the existence of two additional intermediate axes of 𝑢 and 𝑣, which correspond to 𝑈 = 𝑋 ⊕ 𝑌 and 𝑊 = 𝑌 ⊕ 𝑍, respectively. Specifically, the 𝑢 axis can differentiate vertices {1,4,5,8} from vertices {2,3,6,7}, and the 𝑤 axis can differentiate vertices {1,2,7,8} from vertices {3,4,5,6} (Supplementary Fig. 12A&C). Consistent with this prediction, the SVM analysis showed significantly higher classification accuracies for these vertex sets compared to the baseline, confirming the existence of these two intermediate 𝑢 and 𝑤 axes in the high-dimensional neural space (Supplementary Fig. 14A). The characteristics of the 𝑢 and 𝑤 axes were the same as those of the 𝑣 axis: first, the 𝑢 and 𝑤 axes were orthogonal to each other and to other axes (Supplementary Fig. 14B); second, they emerged subsequent to the sensory axes (i.e., 𝑧, 𝑥, and 𝑦 axes) yet prior to the 𝑝 axis (𝐹(2, 697) = 1015.6, 𝑝 < 0.001; for pairwise comparisons, all 𝑝𝑠 < 0.001, Bonferroni corrected) (Fig. 3F, right; also see Supplementary Fig. 14C); third, the cubic XOR problem (i.e., the classification of contour orientations) can be addressed using either the 𝑢 axis or the 𝑤 axis as the intermediate axis (Supplementary Fig. 12). In addition, these 7 axes were present in each of three cytochrome oxidase stripes (i.e., thin, thick, and pale) in the V2 area, suggesting that the twist operation is likely a general property of V2 neurons (Supplementary Fig. 16). Taken together, the dimensionality of the perceptual manifold was at least 7, much higher than that of the sensory manifold (i.e., 3).

An intriguing question arises: why was the perceptual manifold embedded in a 7-D space when a 4-D space, constructed by 𝑥, 𝑦, 𝑧, and 𝑝 axes, is sufficient to satisfy the task demand of classifying contour orientations? One possibility is that the availability of multiple alternative pathways to construct the task-relevant 𝑝 axis enhances the robustness for the classification. Alternatively, the perceptual manifold may not be task-specific; rather, the classification of contour orientations could be just one of its many possible applications. In fact, for the stimulus space with 8 vertices, there are 2^8^ = 256 possible classifications. Some are linearly separable in the stimulus space, such as vertex {1} versus {2,3,4,5,6,7,8} or vertices {1,5} versus {2,3,4,6,7,8} (Fig. 5A), while others are not, such as {1,3} versus {2,4,5,6,7,8} (Fig. 5B). In total, in the stimulus space, 104 classifications are linearly separable, and 152 are not (for a full list, see Supplementary Fig. 17 and Supplementary Table. 1). Critically, all linearly inseparable classifications in the stimulus space become linearly separable in the 7-D space. For example, the vertex pair of {1,3} versus {2,4,5,6,7,8} becomes linearly separable in the 𝑦-𝑧-𝑝 subspace (Fig. 5C). In total, there are 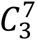 = 35 3-D subspaces embedded in the 7-D perceptual space (Supplementary Fig. 18), and each of the 152 linearly inseparable classifications becomes linearly separable in at least one of these 35 subspaces (Supplementary Table. 2). That is, every possible classification in the stimulus space is linearly separable in this 7-D perceptual manifold.

**Figure 5:**
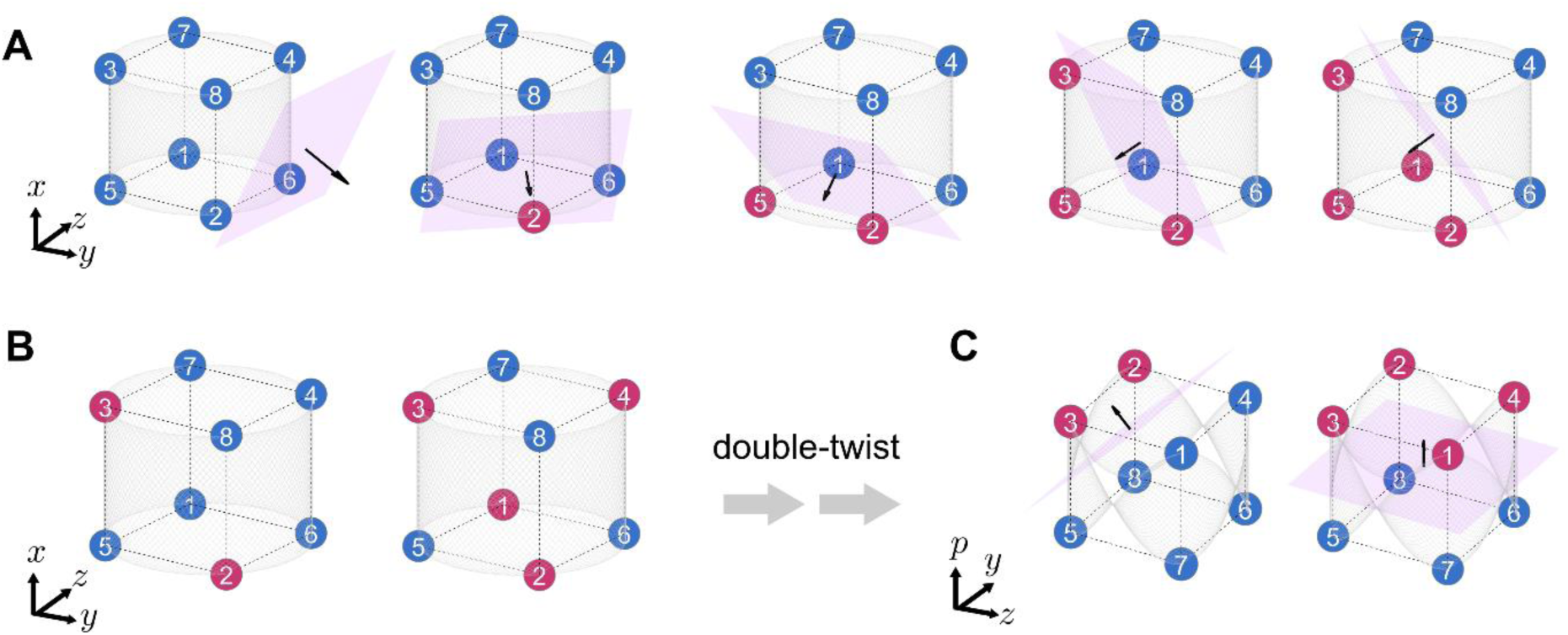
Linearly separable and inseparable classifications in the stimulus space. (A) Examples of linearly separable but non-axial classifications in the stimulus space. The classification separates a subset of vertices (with the set size of 0, 1, 2, 3, and 4, red) from the rest (blue). (B) Examples of linearly inseparable classifications in the stimulus space, including the classification of contour orientations (right). (C) The classifications in panel B become linearly separable in new subspaces through twist operations.

Therefore, the perceptual manifold is unlikely for decision-making or action; rather, it may provide a reservoir of all possible candidates (in our case, solutions for all possible classifications) for higher-order cognitive processes. Consistent with this conjecture, after excluding task-relevant neurons that showed high sensitivity to contour orientations from the V2 neuron population, the remaining neurons retained the ability to classify contour orientations at the population level (Supplementary Fig. 7).

### The necessity and sufficiency of NMS neurons in dimension expansion

The aforementioned analyses showed the important role of NMS neurons in expanding dimensions of representational spaces through twist operations. Here, we further examined the necessity and sufficiency of NMS neurons in dimension expansion. To do this, we first compared NMS neurons with pure selectivity neurons in carrying out all possible classifications in the stimulus space. We generated synthetic neurons exclusively tuned to one of the three stimulus features (Supplementary Fig. 21) based on real recorded data in V2 to simulate neurons with pure selectivity (Rigotti et al., 2013). For example, Fig. 6 A & B show a typical NMS neuron in the V2 area responding differently to HV and CA and showing no sensitivity to OI (top), and a typical synthetic neuron with pure selectivity to OI (bottom).

**Figure 6:**
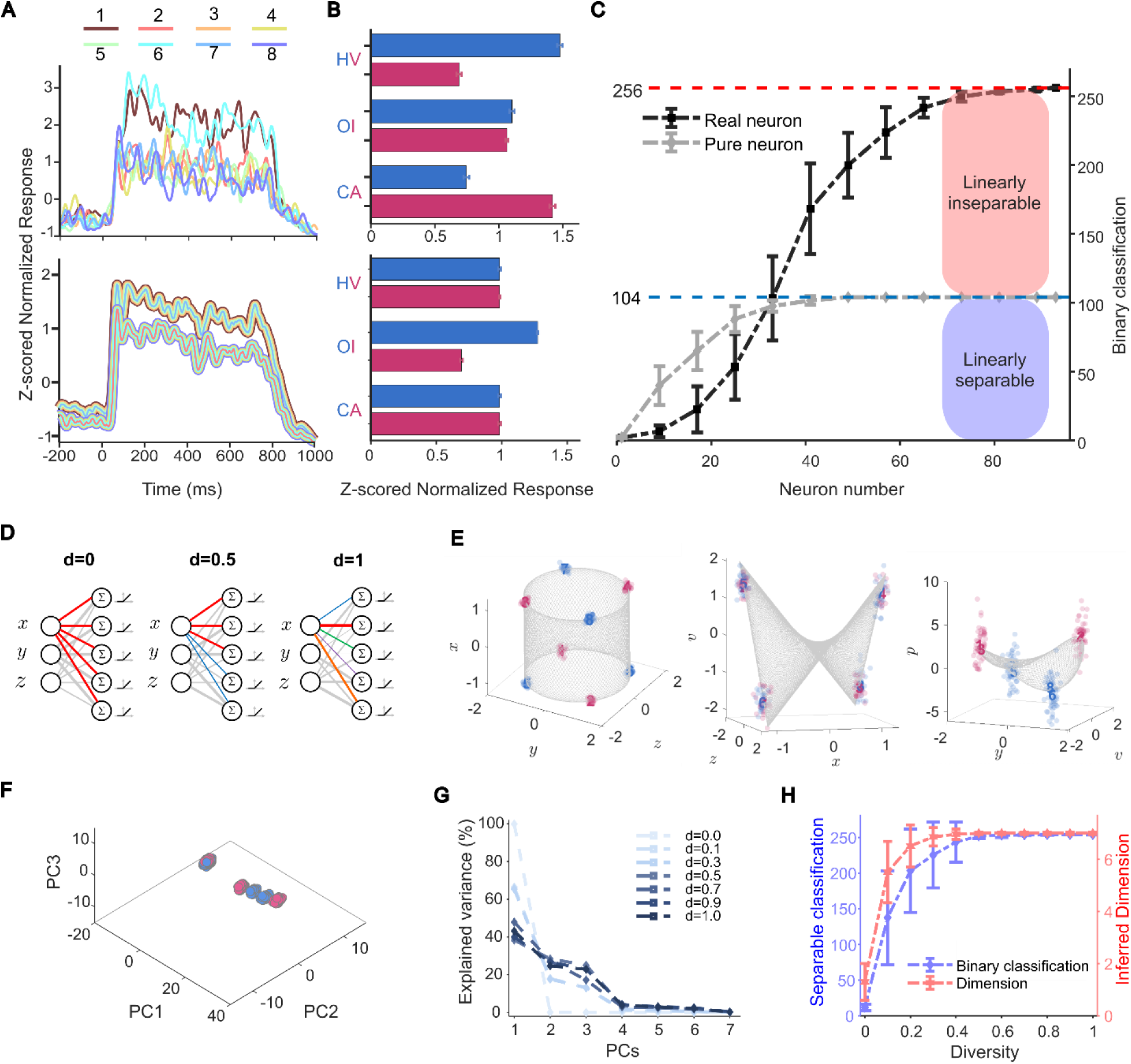
Necessity of NMS neurons and heterogeneous connectivity structure. (A) Time courses of an NMS neuron (top) and a synthetic neuron with pure selectivity (bottom). Colors and numbers denote different stimuli and their associated neural responses. The time course of the synthetic neuron was generated by modifying the time course of the NMS neuron shown above. (B) Average activity of the NMS neuron (top) and the synthetic neuron (bottom) from 200ms to 750ms after stimulus onset. Colors denote different conditions of a feature, and error bars denote the standard error of mean. (C) The number of successful linear classifications increased as a function of neuron population size for both pure selectivity neurons (gray) and NMS neurons (dark). The gray curve plateaus at 104, the total number of linearly separable classifications in the stimulus space, whereas the dark curve converges to 256, including both 152 linearly inseparable classifications and 104 linearly separable ones. Real neuron: neurons recorded in the V2; Pure neuron: synthetic neuron with pure selectivity; error bar: standard deviation. (D) Schematic illustration of a two-layer feedforward neural network. Each output neuron of the network receives the inputs encoding three stimulus features (HV, OI and CA), indexed by 𝑥, 𝑦, and 𝑧, respectively, from the input neurons. The connectivity weights are randomly sampled from a multivariate Gaussian distribution, with the thickness of lines indicting the magnitudes of the weights. Connectivity patterns from feature 𝑥 to the output neurons are highlighted by colors for display purposes. The response of output neurons is achieved through linear summation and ReLU activation. Networks with three levels of diversities in connectivity patterns are shown from left to right. (E) Visualization of neural manifolds in 3-D subspaces when 𝑑 = 1 from a typical simulation. Left: Sensory manifold that corresponds to the stimulus space; Middle: Intermediate manifold after one twist operation; Right: The subspace where linearly classification of contour orientations is achieved. Each point denotes the neural state of a stimulus, with red and blue colors representing the two contour orientations, respectively. (F) Visualization of the neural manifold in a 3-D subspace when 𝑑 = 0 from a typical simulation. The subspace is constructed by the first three PCs of neurons’ activation. (G) Variance explained by PCs of neurons’ activation with different level of diversity in connectivity patterns from a typical simulation for each 𝑑 value. When 𝑑 = 0, indexed by the dashed line with the lightest blue, the variance is nearly explained by the first PC alone. When 𝑑 = 1, indexed by the dashed line with the darkest blue, the curve of variance explained by each PC is flattened. (H) The dimensionality and the number of successful classifications as a function of heterogeneity in response profiles from 100 simulations for each 𝑑 value. Left axis (blue): the number of successful classifications. Right axis (red): inferred dimensionality of neural manifolds constructed by the network (for details, see Methods). Error bar: standard deviation.

To evaluate the classification performance for 152 linearly inseparable problems (Supplementary Table. 1) and 104 linearly separable problems (Supplementary Table. 2), we used a population-increment procedure (Rigotti et al., 2013), where the population size was progressively increased from a single neuron to the entire set of 93 neurons, adding one randomly selected neuron each time. During each iteration, we trained SVMs with neural activities for all possible classifications, and a classification accuracy threshold of 75% was set as the criterion for successful classification (for details, see Methods). Figure 6C shows the number of successful linear classifications as a function of neuron population size. With NMS neurons, we succeeded in all possible classifications (256 in total), for both linearly separable and inseparable problems (Fig. 6C, black curve), when the number of NMS neurons exceeded 81. In contrast, using the synthetic neurons with pure selectivity, the total number of successful classifications plateaued at 104 (Fig. 6C, gray curve) when the population size exceeded 49. That is, additional increase in neuron population size did not further improve classification performance.

Critically, close examination of the problems successfully classified by pure selectivity neurons revealed that they were all linearly separable in the stimulus space (Supplementary Table. 1), and none came from the set of linear inseparable problems (Supplementary Table. 2). That is, neurons with pure selectivity can only address linearly separable problems, as they alone cannot expand the sensory manifold to a higher dimensionality. Taken together, this finding suggests that regardless of neuron population size, NMS neurons are necessary in expanding the dimensionality of neural manifolds, hereby transforming the sensory manifold into the perceptual manifold.

The finding that at least 81 neurons were needed for forming the 7-D perceptual manifold, as shown in Fig. 6C, highlights the importance of population-level activity in dimension expansion. Indeed, previous studies have shown that neurons’ diverse response play an important role in computational capacity (Rigotti et al., 2010; Kriegeskorte et al., 2021; Gast et al., 2024).

To quantify how diversity in response profiles of NMS neurons influences dimension expansion of the representational space, we built a two-layer feedforward neural network tasked with processing the stimuli used in the macaques’ experiment (for details on the network, see Methods). In this network, each output neuron receives the combination of all three stimulus features (i.e., HV, OI and CA) from the input neurons and employs a nonlinear activation function (i.e., ReLU), with connectivity weights independently sampled from a multivariate Gaussian distribution. As a result, all output neurons in this neural network demonstrate a response profile of nonlinear mixed selectivity.

In this network, the response profiles of NMS neurons are controlled by a parameter 𝑑, which denotes the degree of diversity in connection patterns between the two layers (Fig. 6D). This diversity ranges from identical patterns (𝑑 = 0) to completely uncorrelated patterns (𝑑 = 1) (see Methods). When 𝑑 = 1, each NMS neuron generates a distinct response because the connection pattern from the input neurons is unique (Fig. 6D, right), and therefore the matrix of connectivity weights is full rank. SVM analysis, similar to that performed on the macaques’ data, was carried out to identify the 7-D perceptual manifold. For visualization, neurons’ activations are projected into 3-D subspaces (Fig. 6E), where each dot denotes the neural state of a stimulus, with red and blue colors representing the two contour orientations, respectively. Within this 7-D perceptual manifold, we can identify the sensory manifold embedded in a 3-D subspace constructed by axes corresponding to the three stimulus features (Fig. 6E, left), the intermediate manifold in a 3-D subspace with a new axis 𝑣 after one twist operation (Fig. 6E, middle), and the subspace achieved after the second twist operation where linearly separating contour orientations becomes possible (Fig. 6E, right). In addition, continuous stimuli that spanned the entire sheared configuration ring (Fig. 1C, right) produced similar results (Supplementary Fig. 22). In sum, the 7-D perceptual manifold constructed by the network of NMS neurons with random connectivity patterns (i.e., 𝑑 = 1) is comparable to the 7-D perceptual manifold identified in the macaque’s V2 (Fig. 4D).

In contrast, when 𝑑 = 0, all NMS neurons have the same inputs and thus generate identical responses. Accordingly, the matrix of connectivity weights in the network is rank 1 (or 0 if all weights are 0), resulting in low dimensionality of the neural manifold (inferred dimension = 1.3, std. = 0.70, see Methods). This low dimensionality is also revealed is demonstrated by PCA analysis of the variance in neuron activation (Fig. 6G). When 𝑑 = 0, the first PC explained 99.84% of the total variance, leaving nearly no variance for the remaining PCs. As a result, the neural states of the stimuli were confined to an approximately 1-D space (Fig. 6F). In contrast, when 𝑑 = 1, the first 6 PCs (99.78%) were required to explain the same amount of variance as the first PC when 𝑑 = 0. Thus, when 𝑑 = 1, the neural states of the stimuli were dispersed into a higher dimensional neural space. Therefore, the neural manifold constructed by the network with no diversity (i.e., 𝑑 = 0) suffers a severe deficit in performing either linear or nonlinear classification (the number of linearly separable problems successful addressed: 9.16, or 8.8% of the whole set, std. = 2.89; the number of linearly inseparable problems successful addressed: 2.74, or 1.8%, std. = 2.23). These findings suggest that networks consisting of NMS neurons with an identical response profile have limited computational capacity and thus hardly encodes sufficient information, even when the response profile exhibits nonlinear mixed selectivity.

To systematically investigate how diversity in the response profiles of NMS neurons influences the dimensionality of representational spaces, we constructed a series of neural networks with different parameters 𝑑, and then measured the dimensionality and the classification performances (see Methods). We found that as the diversity in response profiles increased, the dimensionality increased monotonically (Fig. 6H, red curve). Interestingly, the network did not need to possess completely diversity to form the perceptual manifold. With 𝑑 > 0.5, the dimensionality reliably expanded to 7 (see Methods and Supplementary Methods M.5). In parallel, the number of successful classifications increased monotonically, finally capable of successfully carrying out all possible classifications (i.e., 256) once the dimensionality reliably reached 7 (Fig. 6H, blue curve). Note that the linearly inseparable problems were resolved in parallel with the linearly separable ones (Supplementary Fig. 23).

On the other hand, heterogeneity in response profiles alone seems insufficient, as neural networks consisting of pure selectivity neurons with identical parameter 𝑑 were only capable of addressing linearly separable problems (Supplementary Fig. 24). Taken together, the synergy between twist operations on input feature vectors by NMS neurons and the heterogeneous response profiles among NMS neurons is critical, which optimally leverages neural networks to construct a more complex, higher-dimensional neural space.

## Discussion

In this study, we investigated how macaques’ V2 neurons solve linearly inseparable problems present in the physical world through the lens of neural geometry. By analyzing the neural geometry embedded in the high-dimensional neural space formed by V2 neurons’ collective activities, we identified two related but distinct neural manifolds: the sensory and perceptual manifolds. The sensory manifold, embedded in a 3-D subspace defined by the stimulus features, faithfully represented the raw sensory input where contour orientations remained linearly inseparable. However, through a series of geometric transformation equivalent to twist operations, this 3-D sensory manifold evolved into a 7-D perceptual manifold with four additional axes, allowing for the linear separability of contour orientations. Critically, this geometric transformation was achieved through the synergy between twist operations on input feature vectors by NMS neurons at the individual neuron level and the heterogeneous response profiles among NMS neurons at the population level. In summary, our study offers mechanistic insights into how biological neural networks expand the dimensions of representational space, transitioning from the sensory to the perceptual manifold, thereby advancing our understanding of how information progresses from sensation to perception.

Previous studies on neural geometry have shown that neural manifolds can faithfully represent both stimulus (Xie et al., 2022; Gardner et al., 2022; Bao et al., 2020) and action spaces (Churchland et al., 2012; Gallego et al., 2017; Perich et al., 2020). In line with these findings, our study identified a neural manifold embedded in a 3-D subspace defined by three mutually orthogonal axes corresponding to the HV, OI, and CA features of the MIC stimuli. Along with the finding that the size of this manifold was found to correlate with the intensity of motion coherence, this manifold reflects the raw sensory input (i.e., the stimulus space) and is therefore termed the sensory manifold. Importantly, our theoretical predictions, derived from the geometric transformation of input feature vectors by NMS neurons, led to the identification of four additional axes that encode features absent from the physical stimuli. Specifically, one of these axes, resulting from double twist operations on the three feature axes of the sensory manifold, encoded the perceived orientations of illusory contours, allowing for linear separability of contour orientations that were not linearly separable in the sensory manifold. Interestingly, this manifold was not specific to the task at hand, as it potentially accomplishes all 256 possible classifications present in the stimulus space. Therefore, this manifold likely functions as an intermediary between the sensory manifold and those associated with decision-making or action, hence its designation as the perceptual manifold. Note that the perceptual manifold observed in the V2 does not necessarily originate and terminate within the V2. It likely inherits characteristics from feedforward inputs from the V1 and is further shaped by feedback from downstream cortical regions such as the V3 and V4. Future research employing simultaneous recordings across multiple areas is essential to elucidate the transformation from the sensory to the perceptual manifold.

The creation of these perception-related axes was attributed to NMS neurons, which have the unique ability to geometrically transform input feature vectors. This transformation is equivalent to twist operations, thus expanding the dimensions of representational spaces (Fusi et al., 2016; Rigotti et al., 2013). In contrast, neurons exclusively selective for a single feature or those exhibiting a linear combination of selectivity for multiple features are unable to change dimensionality. Therefore, NMS neurons appear to be necessary for interpreting sensory inputs into perceptual experiences by generating latent variables from intermediate axes without direct semantic descriptions (Johnston et al., 2020; Jazayeri & Ostojic, 2021). However, the mere presence of NMS neurons is not sufficient; their functional efficacy depends on the heterogeneity of their response profiles at the population level (Langdon et al., 2023; Kriegeskorte et al., 2021). Through simulations of neural networks consisting of NMS neurons, we found that homogeneous response profiles among NMS neurons limited their capacity of expanding dimensions, thereby constraining the network’s computational power in addressing both linearly separable and inseparable problems. Conversely, increasing the heterogeneity of the response profiles significantly enhanced dimension expansion, effectively transforming linearly inseparable problems into linearly separable ones. In summary, our study reveals the symbiotic relationship between the geometric transformation capability of individual NMS neurons and the heterogeneity in the response profiles at the population level, underscoring the importance of both individual neuron properties and population dynamics in achieving dimension expansion.

The high dimensionality of the perceptual manifold functions as a reservoir of computational solutions, enabling flexible classifications according to downstream task demands. That is, one key function of dimension expansion is apparently to facilitate parallel processing, allowing multiple computations (such as classifications in this study) to occur simultaneously across different dimensions. As these computations are distributed across multiple dimensions in parallel, they become less reliant on conscious control and more automatic. However, this parallel processing comes with a cost, as the number of potential solutions increases exponentially with the number of dimensions. In our study, downstream cortical regions involved in decision-making, which is usually sequential processing, must select the appropriate classification from 256 possibilities to meet the task demands. While the mechanism for effectively navigating these potential solutions based on task demands remains largely unknown, the modulation of NMS neurons’ response profiles through Hebbian (“fire together wire together”) and anti-Hebbian (“out of sync, lose the link”) rules (Caporale & Dan, 2008) might offer insights into reducing dimensionality and thus narrowing the range of potential solutions. In networks governed by Hebbian plasticity, neurons frequently co-activated by similar tasks or stimuli develop more homogeneous response profiles, leading to the formation of specialized modules with relative lower dimensionality (Bullinaria, 2007; Lindsay et al., 2017; Weigand et al., 2017). This idea is supported by recent findings showing that sequences of contextually related images (e.g., natural video) are represented in a neural space with lower dimensionality, evidenced by straighter neural population trajectories, compared to sequences of contextually unrelated stimuli (Hénaff et al., 2021).

Conversely, under anti-Hebbian rule, task-relevant features are disentangled from task-irrelevant ones, which may result in the compression of axes representing these task-irrelevant features, thereby reducing effective dimensionality (Rajan et al., 2010; Ganguli et al., 2012; Farrell et al., 2022). This type of geometric transformation reshapes the representational space to focus on task-relevant features (Bernardi et al., 2020). This conjecture is supported by our findings that although the perceptual manifold can accomplish all 256 classifications, the accuracy along some dimensions (such as CA axis) was lower than others, suggesting that not all features are equally represented in the neural space. Taken together, along the hierarchy of the ventral visual stream, the amount of information encoded in each dimension varies significantly (Stringer et al., 2019), and the relational structure between representations evolves from relatively simple and straightforward to more abstract and complex, reflecting the integration of multiple features and the emergence of high-level perceptual categories (Lin & Kriegeskorte, 2023). This dynamic transformation further orchestrates the progression from perceptual to decision-making manifolds and ultimately to action-oriented manifolds, through intensive interplay of top-down and bottom-up processing, enabling us to act upon the physical world in response to stimuli that have acted upon us.

## Supplemental Information

Supplemental Materials includes 25 figures, 2 tables and 2 videos.

## Acknowledgments

We thank Dr. Xingyu Liu for her valuable discussion and comments, and Dr. Haidong Lu for his supervision on conducting the experiments. This study was funded by Beijing Municipal Science & Technology Commission, Administrative Commission of Zhongguancun Science Park (Z221100002722012), Tsinghua University Guoqiang Institute (2020GQG1016), and Beijing Academy of Artificial Intelligence (BAAI).

## Author contributions

H.M., L.J., T.L. and J.L. developed the theoretical framework. H.M. performed the experiments, collected the data, and compiled the data. H.M., L.J., T.L., and J.L. conceived the data analyses. H.M. and L.J. performed the data analyses. L.J. developed the double-twist model. T.L. performed the network simulation. H.M., L.J., and J.L. wrote the paper.

## STAR★METHODS

### KEY RESOURCES TABLE

| REAGENT or RESOURCE | SOURCE | IDENTIFIER |
| --- | --- | --- |
| **Experimental Models: Organisms/Strains** |  |  |
| Rhesus macaque (*Macaca mulatta*) | Suzhou Xishan<br>Zhongke animal<br>Company, Ltd; | http://xsdw.bioon.com.cn/ |
| **Deposited data and code** |  |  |
| Codes and dataset | This paper | https://data.mendeley.com/datasets/vrj3645vgz/2 |
| **Software and Algorithms** |  |  |
| MATLAB | MathWorks | R2017b & R2022b |
| Jupyter | Project Jupyter |  |
| **Other** |  |  |
| 32channel-Array | Blackrock<br>Microsystems, LLC; | https://www.blackrockmicro.com/ |
| 64channel Multichannel Neural Recording<br>(AlphaLab SNR) | Alpha Omega; | https://www.alphaomega-eng.com/ |
| ViSaGe | Cambridge Research<br>System Ltd; | https://www.crsLtd.com/ |

## RESOURCE AVAILABILITY

### Data and materials availability

The data and analysis code are available at Mendeley Data, https://doi.org/10.17632/vrj3645vgz.2. All data information that supports the findings of this study are available from the corresponding author on reasonable request.

### Materials and Methods

Four hemispheres from two adult male macaque monkeys (Macaca mulatta) were used in this study. All procedures were performed in accordance with the National Institutes of Health Guidelines and were approved by the Institutional Animal Care and Use Committee of the Beijing Normal University (protocol number: IACUC(BNU)-NKCNL2013-13).

### Visual stimuli, behavioral tasks, and recordings

The data used in this study was from our previous work, and for more details on stimuli, tasks, and recordings, see Ma et al, 2021. Stimuli were generated with ViSaGe using MATLAB scripts and presented on a 21-inch CRT display. MIC stimuli had seven levels of dot motion coherence. The size of the MIC was a 4°-diameter circular and the position of MIC was consistent with array population RF.

Two monkeys performed an MIC orientation-discrimination task after headpost and optical imaging guided 32-channel Utah array implant. This task was a two-alternative forced-choice discrimination task and the monkeys were trained to make an eye saccade choice based on MIC orientations. The monkeys made a saccade to the right target if the orientation was tilted to the right of the vertical axis, and vice versa. The monkeys received water reward for correct choices.

The electrophysiological recording system is AlphaLab SnR 64-channel system. Neural signals were sampled at 22 kHz and with an 800 – 7500 Hz bandpass filter. Recordings were performed on multiple days. In this study, we only used single neurons in the unique-unit dataset (Ma et al., 2021). This dataset was generated by excluding potential duplicated units (i.e., similar waveforms or tunings) that were recorded from the same electrodes on different days. Therefore, the neurons in this dataset were either from different electrodes or from the same electrode but had different waveforms or tunings. Additionally, we further refined our selection to include only single neurons from this dataset.

### Data analysis

#### Data preprocessing

For all single neurons selected as previously described, they passed the receptive field (RF) test. Briefly, we used two types of RF mapping stimuli. One is grid-like RF mapping, where a 0.8° square-wave grating is presented at different positions on the grid. We fit the neuronal response in two dimensions using a 2-D Gaussian function; the other is 4° long, 0.2° wide bars presented at different horizontal and vertical positions, for which we use a one-dimensional Gaussian for fitting. A goodness of fit greater than 0.7 is considered as passing the RF test, see Ma et al, 2021. Then, to build a high-dimensional neural space, we identified all V2 neurons that participated in the MIC orientation-discrimination task. In total, we obtained 93 V2 single neurons, with 47 neurons from Monkey S and 46 neurons from Monkey W.

We sorted all trials into 112 conditions to analyze each neuron’s response (2 motion-axis orientation conditions, 4 sheared configuration conditions, 7 coherence levels, and 2 performance outcomes). Subsequently, we calculated each neuron’s trial-averaged response (from -200ms before stimulus onset to 200ms after stimulus offset) with a Gaussian window (a 10ms sliding window with a 2ms step size). We then combined all neuron responses after z-scoring each neuron’s trial average response (Okazawa et al., 2021; Mante et al., 2013). Additionally, we excluded conditions with fewer than 3 trials for some neurons, so we totally got 61 useful conditions. Following these steps, we constructed a data matrix of dimensions 61 (useful conditions) x T (trial time) x 93 (neuron number) from the MIC orientation discrimination task.

#### Support vector machine (SVM)

We used SVM for two purposes: first, to decode categorical information from the neural data, and second, to provide a well-defined vector, which represents a distinct dimension in the neural space. In the analysis, we used the *fitcsvm* function with a linear kernel in MATLAB. We retained the default hyper-parameter values of the function, except for customizing the box constraint. In the neural geometry part, the box constraint was set at 0.001, whereas for the binary classification part, it was set at 1 (for an explanation of box constraints, see Supplementary Methods M.2). Then, the SVMs were trained and tested using data points in the neural space.

Since our population data is composed of multiple sessions with varying trial numbers, we followed previous methods (Mante et al., 2013; Okazawa et al., 2021) and used trial-averaged data (as described in Data Preprocessing). So here we divided the data at different time points into training and testing sets. First, we partitioned the complete time range from -200ms to 1000ms relative to stimulus onset into tiled 12ms-wide time bins. Within each time bin, we randomly selected half of the time steps for training and the other half for testing. To train a single SVM classifier, the selected training data from time bins were pooled together. To generate time courses of classifier accuracy, the classifier was tested within each time bin using the testing data. We also tried another method where, instead of using tiled 12ms-time window, we randomly selected half of the time points from the entire time span as training data and the other half as test data. The classification accuracies were consistent. However, in this way we could not obtain time courses of classification accuracy, so we did not use it in this text.

#### Angle analysis

To calculate the angle subtended between two 𝑛-dimensional unit vectors 𝜷_𝒊_ and 𝜷_𝒋_, we used the following formula:

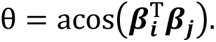

The vectors were from SVM classifications. For the neural geometry and twist model parts, we performed SVM analysis once for each classification. For control analyses, we ran SVM analysis 200 times for each classification (see Supplementary Notes N.6 and Supplementary Fig. 6 A & B). The orthogonality of the angle was tested by examining whether it significantly differed from the angle distribution constructed by randomly selecting two vectors in a 93-dimensional space (see Supplementary Notes N.6 and Supplementary Fig. 6C).

#### Low-pass filter

To smooth the temporal profiles of the neuron activities, we applied customized simple discrete-time RC low-pass filters. Let the temporal profile of a neuron be 𝑥(𝑡). The filtering is applied using a sliding window starting from 𝑁 time-steps before the current moment. In this time window, the filtered temporal profile 𝑥̅(𝑡) is

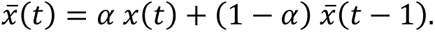

where 𝛼 is the smoothing factor. The factor 𝛼 is computed from the sampling time interval Δ𝑡 and the required cutoff frequency 𝑓_𝑐_ as

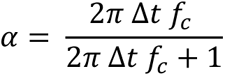

Essentially, the low-pass filter computes the exponentially weighted moving average of the original temporal profile. In our study, Δ𝑡 = 0.002s, 𝑓_𝑐_ = 2Hz and 𝑁 = 5.

#### Three-D visualization

The main axes identified by SVM must be perfectly mutually orthogonal for creating Cartesian coordinate systems. So here we first created an arbitrary full rank matrix 𝑨. Its first few columns were replaced by the identified main axes by SVM. We then applied QR decomposition on matrix 𝑨 to obtain an orthogonal matrix 𝑸. The transformation matrix was 𝑻 = 𝑸^T^(see Supplementary Methods M.3 for details). Then, we used this transformation matrix to linearly transform the original neural space to a new coordinate system where the identified main axes by SVM were the first several axes.

To neatly visualize the neural states in a 3-D subspace, we first applied low-pass filter to the data in the original neural space. We chose cutoff frequency 𝑓_𝑐_ = 2Hz, since we wanted to smooth the curve for better visualization and clearer depiction of the dynamic process. Lowpass filtering was not used in other quantitative calculation. We then used matrix 𝑻 to transform the filtered data into a new high-dimensional coordinate system. Finally, the transformed data were projected into a 3-D subspace constructed by the main axes identified for visualization.

#### Double-twist model

The double-twist model transformed the continuous cylindrical stimulus manifold depicted in Fig. 4C (left) to a continuous perceptual manifold embedded in a 7-D space. Its projections into 3-D subspaces were shown in gray in Fig. 4C (middle & right). The perceptual manifold arose from the correspondence between the XOR operator and the arithmetic product. Let the true value be represented by -1, and the false value by 1. The truth table of the XOR operator aligns with that of the arithmetic product (see Supplementary Fig. 25). That is,

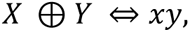

where 𝑋 and 𝑌 are Boolean variables and 𝑥 and 𝑦 are the coordinates on the 𝑥 and 𝑦 axes, respectively. Therefore, after two twist operation, the 7-D coordinates of a point in the perceptual manifold corresponding to a point [𝑥, 𝑦, 𝑧] in the stimulus manifold are [𝑥, 𝑦, 𝑧, 𝑥𝑦, 𝑦𝑧, 𝑥𝑧, 𝑥𝑦𝑧].

The derived perceptual manifold was fit to the neural data using affine transformation for visualization. Because the neural data contained noise, we first applied a low-pass filter (cutoff frequency: 2Hz), and then calculated the steady-state averages (from 300ms to 500ms relative to stimulus onset) of neural activities for the 8 stimuli to determine 8 centers. These centers were then projected into various 3-D subspaces (Fig. 4). In each subspace, we located the neural states for the 8 stimuli based on the derived perceptual manifold. Using the least-square method, we obtained the transformation matrix 𝑭. The mean of the residuals was represented as a vector 𝒆. For any point 𝒙 on the 3-D projections of the derived perceptual manifold, we applied the transformation

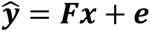

to fit the model to the data (see Supplementary Methods M.4 for detailed formulation). The goodness of fit was measured by R^2^.

#### Binary classification

Binary classification entailed sorting stimulus conditions into two classes based on all possible classification rules. With 8 stimulus conditions, we had a total of 256 classification rules. For each classification, the criterion for linear separability was set at 75% accuracy within each stimulus condition. Each neuron pool was randomly selected from the entire set of 93 neurons, one at a time, and we then conducted SVM analysis 10 times for each neuron pool (see Supplementary Methods M.5).

To infer the dimensionality from binary classifications, we employed the approach developed by Rigotti et al, 2013. Briefly, we calculated the ratio of the actual number of linearly separable classifications and the theoretical number of binary classifications for each condition number 𝑛 (𝑛 ∈ {2, …, 8}). The last 𝑛 whose ratio > 0.8 was selected as the dimensionality (see Supplementary Methods M.6).

#### Connectivity patterns of neural networks

We created a simple two-layer feedforward neural network; the first layer contained 3 units representing the three feature dimensions, respectively, and the second layer contained 93 nonlinear mixed selectivity neurons. Each neuron received the input signals of all the three feature dimensions (mixed): 𝑥(HV), 𝑦(OI) and 𝑧(CA), and employed a ReLU activation function (nonlinearity). The input signals represented the eight corners of the cube in the 𝑥-𝑦-𝑧 stimulus space, each corresponding to a specific stimulus. The activity 𝑟_𝑖_ of the 𝑖th neuron was defined as:

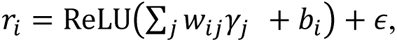

where 𝛾_𝑗_ ∈ {𝑥, 𝑦, 𝑧} is the input signal of the 𝑗th stimulus feature with 𝑤_𝑖𝑗_ as the weight, 𝑏_𝑖_ is a random bias sampled from the uniform distribution 𝒰(0,1), and 𝜖 is a noise term drawn from the Gaussian distribution 𝒩(0, 1/3).

The weights 𝑤_𝑖𝑗_ were randomly sampled from standard Gaussian distributions. The covariance between the weights controlled the structure of the network and thus determined the heterogeneity of the output activities. Hence, we used a diversity parameter 𝑑 ∈ [0,1] to define the 93×93-D covariance matrix 𝐾 for sampling the weights of feature 𝑥:

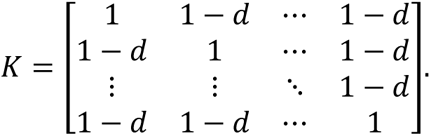

The covariance matrix 𝐾 was fed into the *multivariate_normal* function in Python’s *numpy.random* package to generate the weights. The procedure for generating weights of features 𝑦 and 𝑧 was identical.

We ran multiple simulations on the network. In each simulation, input stimuli were repeated 100 times to allow the added random noise for generating point clusters. We applied the same analyses for the network’s output activities as we did for the neural data. One hundred simulations were conducted for calculating the dimensionality of the neural geometry.

