## Supplemental files for "From Sensory to Perceptual Manifolds: The Twist of Neural Geometry"

### Supplementary Notes

In this Supplementary Notes, we present extended data of empirical results and theoretical derivations.

#### N.1. MIC stimuli and stimuli space

The standard MIC stimuli consist of two main parameters. Supplementary Fig. 1A represents the orientation of the axis of oppositely moving dots, while Supplementary Fig. 1B represents the sheared angle of the neighboring dots moving in the same direction (see Fig. 1B in the Result section). Our work focused on four conditions (represented by black dots in Supplementary Fig. 1B), while other works (referred to as KEG and KEO stimuli, Marcar et al., 2000; Mysore et al., 2006) explored the additional conditions, essentially sampling from the sheared configuration space.

For the MIC orientation discrimination task, we utilized the standard MIC stimuli and added 7 levels of coherence to the moving dots (Ma et al., 2021). Specifically, we selected horizontal-vertical conditions (±15° compared to horizontal or vertical orientation) for the motion-axis orientation and used the four conditions mentioned earlier for the sheared angle. This resulted in a total of 8 stimuli per coherence level. By doing this, we could project these two independent parameters (each having one intrinsic dimension) into 3 dimensions in Euclidean space under Cartesian coordinates.

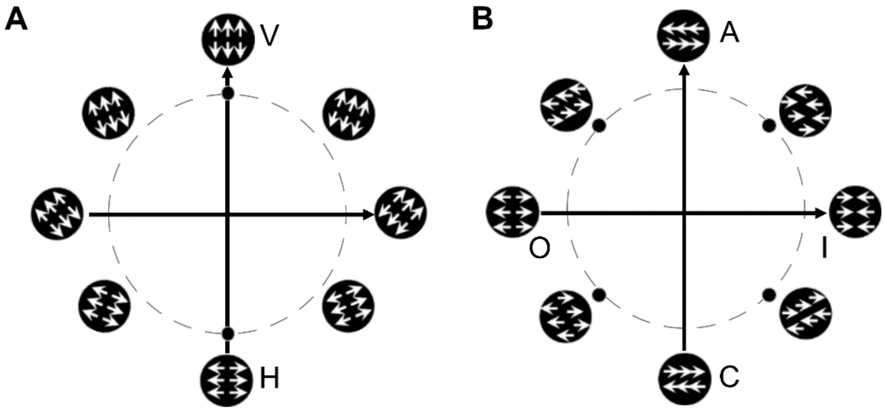


**Supplementary Figure 1. Whole stimuli space.**

The whole stimuli space for MIC. MIC stimuli were constituted by two intrinsic dimensions. Panel A represents the orientation of axis of oppositely moving dots, and we chose 2 conditions (black dots) for further analysis. Panel B represents sheared angle of the neighboring dots moving in the same direction, and we used 4 conditions (black dots) in this sheared configuration space. In this way, we constructed three stimuli dimensions HV, OI and CA to describe the stimulus space of the discrimination task.

### N.2. Sheared configurations of stimuli

To further explain what sheared configurations are, we illustrated how they determine the dot movement of the stimuli and how they were parameterized as a complete space. In the ideal stimuli, the dots move coherently in the two halves of the viewing area in opposite directions. The sheared configurations specify the relative locations of neighboring dots where the coherent movement experiences abrupt change, that is, between the two oppositely moving halves of the stimuli. In other words, it is where the moving dots appear or disappear. Supplementary Fig. 2 A & B depict two examples of sheared configurations for when the motion axis is horizontal. Only half of the viewing area is used in the illustration. The sheared configurations were parameterized by an angle $\theta$ with respect to the motion axis. Due to the coherent motion of the dots, the sheared configuration between any two neighboring dots remains constant through the dot presentation in the viewing area. The two $\theta$s in Supplementary Fig. 2 A & B conceptually are sampled from a circle (see color coding) in which each point represents a distinct sheared configuration for $\theta \in [0, 2\pi]$ (Supplementary Fig. 2C). The circle lies in a 2-D space which can be spanned by two orthogonal axes, each as the feature of the space. The names of the feature axes derive from the stimuli patterns directly on the axes, respectively.

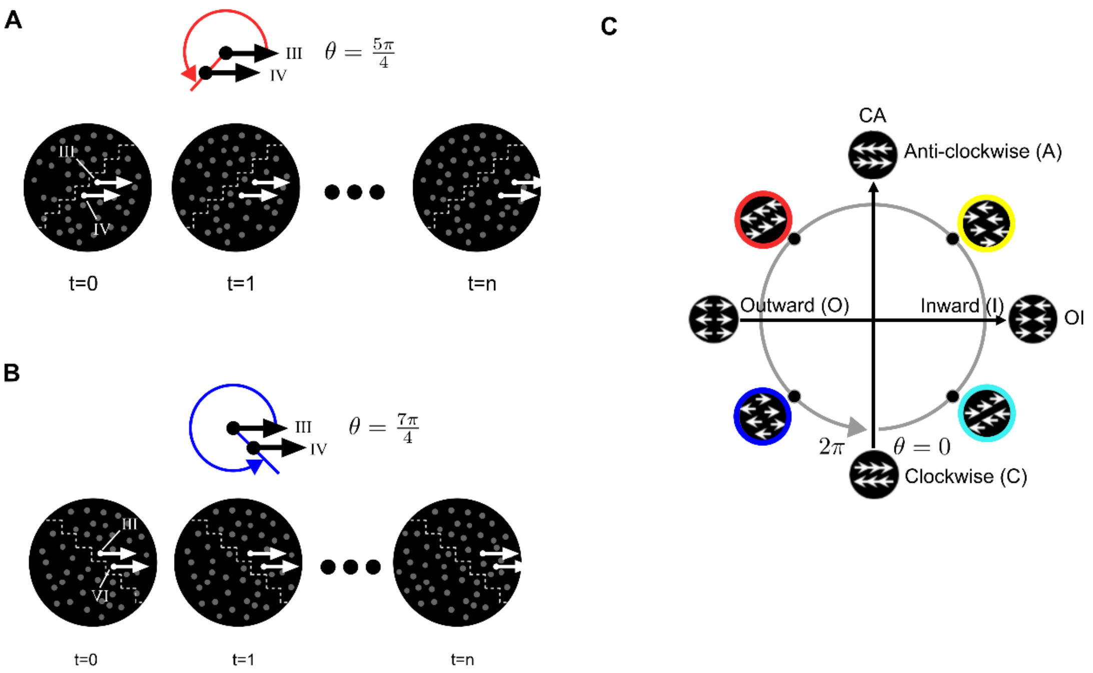


**Supplementary Figure 2. Sheared configurations of stimuli.** (A) An exemplar sheared configuration for neighboring moving dots ($\theta = 5\pi/4$). Because of the coherent motion of the dots, this particular sheared configuration remains constant throughout the dot presentation in the viewing area. (B) Another exemplar sheared configuration ($\theta = 7\pi/4$). (C) The whole space of the sheared configurations. These configurations collectively form a 2-D circle, with $\theta \in [0, 2\pi]$ and $\theta = 0$ positions at the bottom of the circle. Two orthogonal axes extend across the 2-D space where the circle is embedded. According to stimulus patterns on the axes, the axes are therefore named the outward-inward (OI) and clockwise-anticlockwise-like (CA) axes, respectively.

### N.3. Topology of the entire stimulus manifold

The focus of this study enabled us to reduce the stimulus manifold to a cylindrical manifold (Fig. 1D). This is because we used only the horizontal-vertical motion axes in the stimulus set (explained in Supplementary Fig. 1). However, the complete stimulus manifold is more complex; it is a Clifford torus in a 4-D space whose topology is visualized in Supplementary Fig. 3 (left). The cylindrical surface is a 3-D projection of the 4-D torus (Supplementary Fig. 3, right).

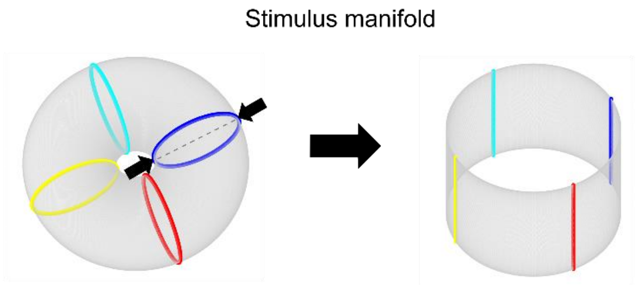


**Supplementary Figure 3. Topology of stimulus manifold.** The entire stimulus manifold takes the form of a Clifford torus in a 4-D space (Cueva et al., 2021). Left: For visualization, the topology of the stimulus manifold is illustrated in a 3-D form, where each colored circle corresponds to a subspace of motion axes. Distinct motion axes are represented by points on these circles, with the color codes inherited from the sheared configurations depicted in Supplementary Fig. 2C: Given our study's focus on just two conditions in the motion axis subspace, namely horizontal and vertical, we can project the 4-D Clifford torus along an unrelated axis (as shown by the dashed line) within the motion axis subspace. This projection transforms the torus into a cylinder embedded in a 3-D space, thereby converting the colored circles into line segments (Fig. 1D).

### N.4. Receptive Field Relative with Stimulus and Stimulus Selective Property of Each Neuron.

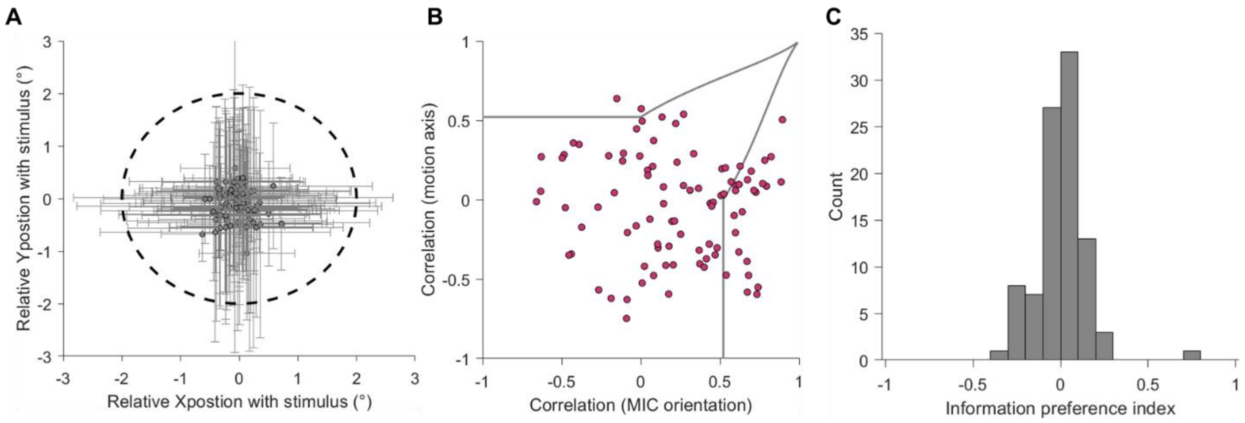


**Supplementary Figure 4. Neurons' receptive fields and selectivity.** (A) Spatial alignment of neurons' receptive fields (RFs, the horizontal and vertical lines represent the range of the RFs centered by a circle point for the neurons.) with the size of the stimuli delineated by a dotted circle. The size of the stimuli was slightly larger than that of neurons' RFs (2.50±0.90°). (B) Neurons' selectivity based on a partial correlation analysis (Marcar et al., 2000; Priebe et al., 2003; Smith et al., 2005). The data used here was from testing MIC basic tuning under passive fixation (Ma et al., 2021). Following the methods of previous researchers (Marcar et al.,

2000), we had two sets of MIC stimuli, each set contained 12 contour orientation conditions, differing in the angle between contour orientation and motion axis. The abscissa represents the correlation coefficient between neural responses for these two sets according to MIC orientation. The ordinate represents the correlation coefficient between neural responses for these two sets according to motion axis. Neurons closer to the top-left showed greater responsiveness to motion information (motion axis), while those nearer the bottom-right were more influenced by MIC information. (C) Neurons' selectivity using the Epoch-preference index method (Elsayed et al., 2016). Similar to panel B, this index was used to compare neural responses driven by MIC orientation and motion axis. Positive values indicate that a neuron is more selective for the motion axis, while negative values indicate that a neuron is more selective for MIC orientation. The distribution of this index did not significantly deviate towards bimodality (Hartigan's dip test; dip statistic = 0.023; $p$ = 0.99), indicating that neurons were approximately equally responsive to both motion and MIC information.

### N.5. Population Average Response and Principal Component Analysis (PCA)

To measure the neural data's ability to differentiate four types of classifications (HV, OI, CA, and LR), we employed traditional methods, including population average response analysis and PCA. In population average response analysis (Supplementary Fig. 5A), we first z-scored the trial average response of each neuron. For one type of classification (e.g., H versus V in the HV classification), we selected higher average responses and represented them as warm color curves, while lower average responses were represented as cold color curves. By aligning all neuron responses based on higher or lower responses, the difference in response between warm and cold color curves indicated the neural data's ability to differentiate the HV conditions.

For the PCA analysis (Supplementary Fig. 5B), we used the high-dimensional neural data and considered each neuron as one dimension. Through the PCA analysis (Matlab2017b toolbox), we obtained 93 orthogonal principal components (PCs). We then projected the population neural responses from the time interval of 200–700ms into this orthogonal PCs space. The PCA results revealed that the 8 conditions formed 8 clusters, and more than three PCs had significant loading weights for these clusters. Because PCA is a linear transformation, it allowed us to transform our neural data from a neuron-based representation to a PC-based representation, where each PC was found by variance in the data. However, it is important to note that individual PCs did not have actual meanings and thus lacked direct interpretability. Our data showed that more than one PC contributed to each of the HV, OI, CA, and LR classifications, indicating that the classification directions of the classifications could not be precisely obtained through the PCA analysis. Consequently, we chose the linear SVM method to identify the classification directions for further analyses.

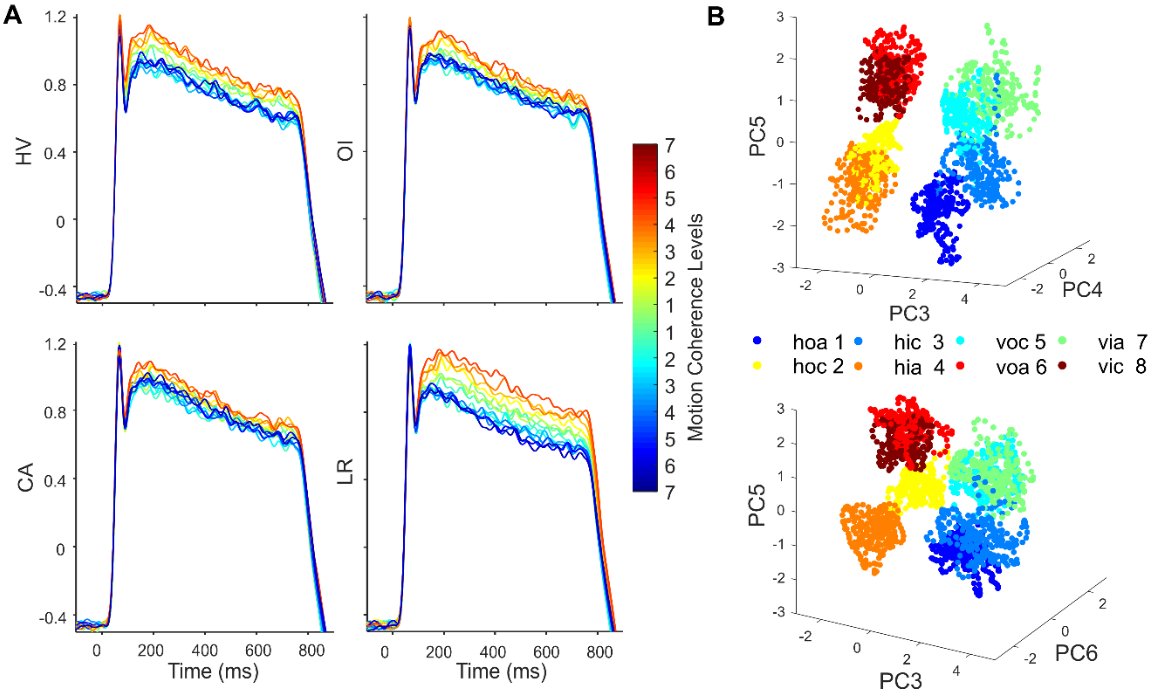


**Supplementary Figure 5. Population related results.**

(A) Z-scored population results aligned by related preference. Four panels represent four types of classifications. Different colors represent the curves for different signal strengths, symmetrically. These panels reflect the ability to measure the difference between two conditions of a category with the traditional average response method. Colors denote levels of motion coherence.

(B) PCA results. The top and bottom panels used different PCs to show 3-D projections. Eight different colors represent 8 conditions. The 8 stimulus conditions formed 8 clusters, and more than 3 PCs had significant loading weights for these clusters.

### N.6. Angle analysis

**Angle distribution.**

For neural geometry in Fig. 2 and Fig. 4, we performed SVM analysis once for each of the HV, OI, CA, and LR classifications and obtained the classification directions. Accordingly, Supplementary Fig. 6A shows the distribution of $\binom{4}{2} = 6$ angles of the 4 different classification directions. For control analysis, in Supplementary Fig. 6 B & C, we ran SVM analysis 200 times for each classification (training and testing sets were directly sampled in the range from -200ms to 1000ms randomly) to obtain 200 directions for this classification. So, we got

$\binom{4}{2} \times 200 \times 200 = 240{,}000$ angles between directions of any two different classifications and $\binom{200}{2} \times 4 = 79{,}600$ angles between directions of a same classification among the 4 different classifications. These two angle distributions are shown in Supplementary Fig. 6B. The angle distribution of different classifications was significantly larger than the angle distribution of a same classification ($p < 0.001$, t-test), with mean values of 92.69 ± 1.94 degrees and 6.82 ± 1.06 degrees, respectively (mean ± standard deviation).

Because the angle distribution between two random vectors is likely more concentrating to 90 degrees as space dimension becomes higher (Widdows & Cohen, 2015), to validate whether the orthogonality observed in Supplementary Fig. 6A was different from randomly selecting two vectors in high-dimensional space, we constructed an angle distribution between two random vectors in the 93-dimensional space (Supplementary Fig. 6C). For this calculation, we followed the method mentioned below (i.e., "Angle between two random high-dimensional vectors"). We found that the resulting distribution (90.02 ± 5.97 degrees, mean ± standard deviation) was significantly different ($p < 0.001$, Kolmogorov-Smirnov test) from the angle distribution of different classifications. The angle distribution of random vectors had a larger standard deviation. Additionally, this wider distribution was similar to the angle distribution obtained after shuffling data labels.

**Angle between two random high-dimensional vectors.**

Theoretically, when two-unit vectors are randomly directed in a $n$-dimensional space and $n$ is large, the subtended angle is close to 90° (Widdows & Cohen, 2015). To demonstrate that our classification directions are indeed orthogonal with each other, we conducted a control analysis by sampling the subtended angles of two randomly directed n-dimensional vectors. We followed the procedure of sampling one randomly directed vector (Muller, 1959), that ensures all the sample vectors are uniformly distributed on the surface of an $(n-1)$-sphere (Knuth, 1973; Poland, 2000). The procedure is as follows:

1. Sample the $n$ components of a vector $\boldsymbol{\beta}_i$ independently from standard normal distribution $\mathcal{N}(0,1)$.
2. Normalize the sampled $\boldsymbol{\beta}_i$ to obtain a unit vector.

We randomly sampled 240,000 pairs of vectors, which is the same number as the angles of different classifications in Supplementary Fig. 6B. For each pair, we calculated the subtended angle between the two vectors, resulting in an empirical angle distribution.

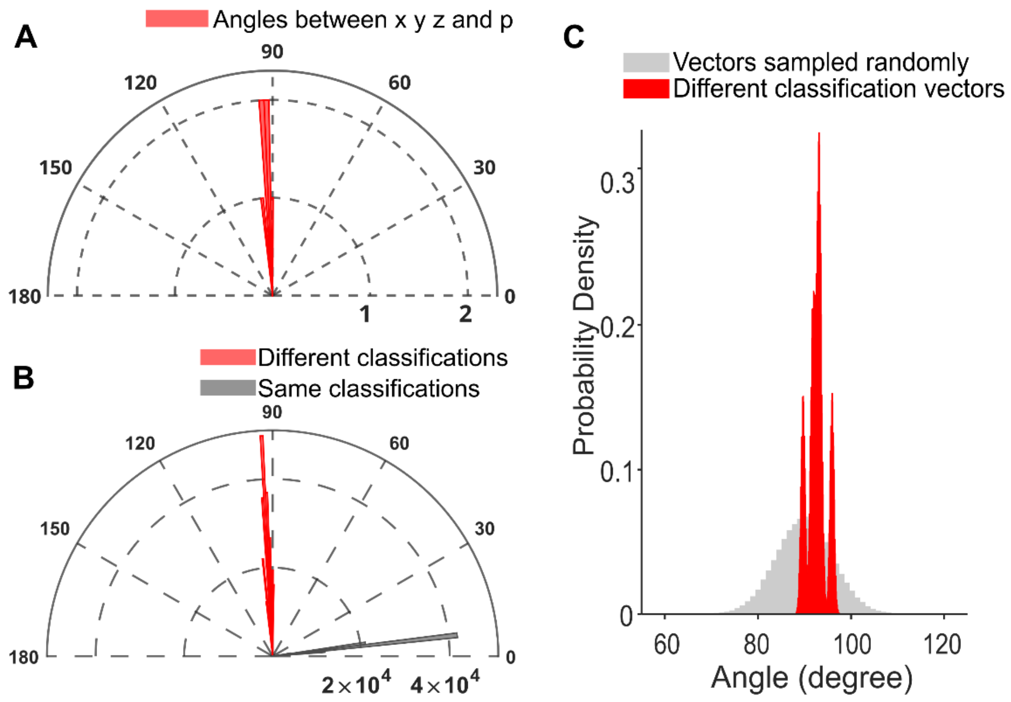


**Supplementary Figure 6. The control analysis of angle distribution.**

(A) Angle distribution of the HV ($x$), OI ($y$), CA ($z$), and LR ($p$) classifications. Angles were calculated for each pair of vectors in the 4 classification directions. The angles concentrated tightly around 90°.

(B) Angle distributions of different classifications versus angle distributions of a same classification obtained from SVM. The distribution of different classifications (red) concentrated tightly around 90°. The distribution of a same classification (grey) concentrated tightly near 0°.

(C) Angle distributions of different classifications versus angle distributions of random vectors in a 93-dimensional space. The distribution of different classifications was significantly more concentrated than the distribution of random vectors.

### N.7. Neural contribution and population coding

To study the characteristics of neural population encoding, we analyzed neural loading weights for each dimension, HV, OI, CA, and LR, and performed SVM analysis while gradually reducing the number of neurons. The results showed that each neuron had mixed selectivity for all dimensions (Supplementary Fig. 7A). For the reducing neuron analysis, after removing neurons with less mixed selectivity, the classifier accuracy did not decrease sharply; instead, the accuracy decreased gradually with smaller number of neurons (Supplementary Fig. 7B).

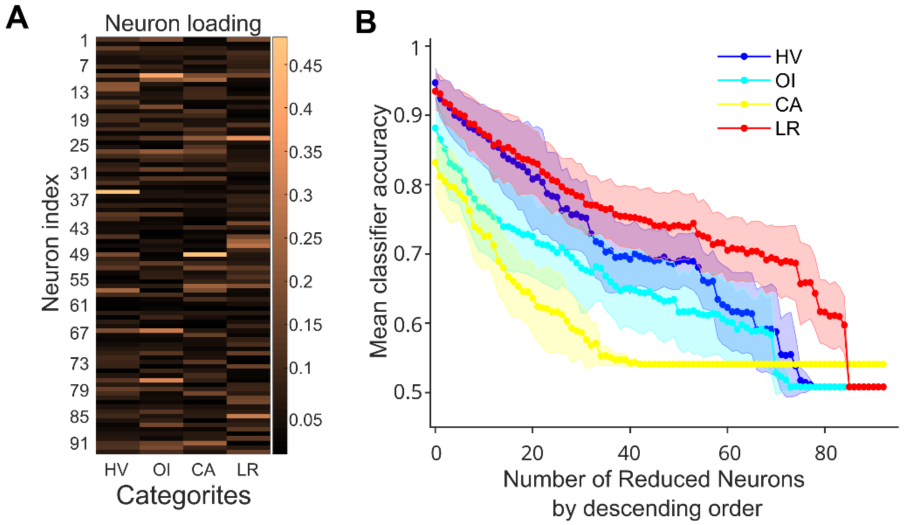


**Supplementary Figure 7. Neuron loading weights and the signature of population coding.**
(A) The matrix of neuron loading weights for the 4 classifications: HV, OI, CA, and LR. Brown color intensity represents the absolute value of loading weights (i.e., the component of a given classification direction on each neuron). As seen, majority of the neurons had bright color, indicating their participation in specifying the classification directions.
(B) Relationship between mean classifier accuracy and the number of neurons. Neurons were removed based on the descending order of the absolute value of their loading weights for each type of classification. Curves with shaded areas represent mean accuracy ± 3 standard deviations (from 200ms to 750ms), and colors represent the 4 types of classifications. Analyses of the loading matrix and the gradual removal of neurons showed that each single neuron contributed to the classifications, and the robustness of classifier accuracy did not decrease sharply upon reducing more selective neurons.

### N.8. Motion coherence on neural manifolds

The size, but not the geometry, of neural manifolds was modulated by the levels of motion coherence. Supplementary Fig. 8A showed cubic vertices on concentric sensory manifolds at different coherence levels, with the highest level of motion coherence showed the largest cubic structure. The similar effect was observed along the LR axis.

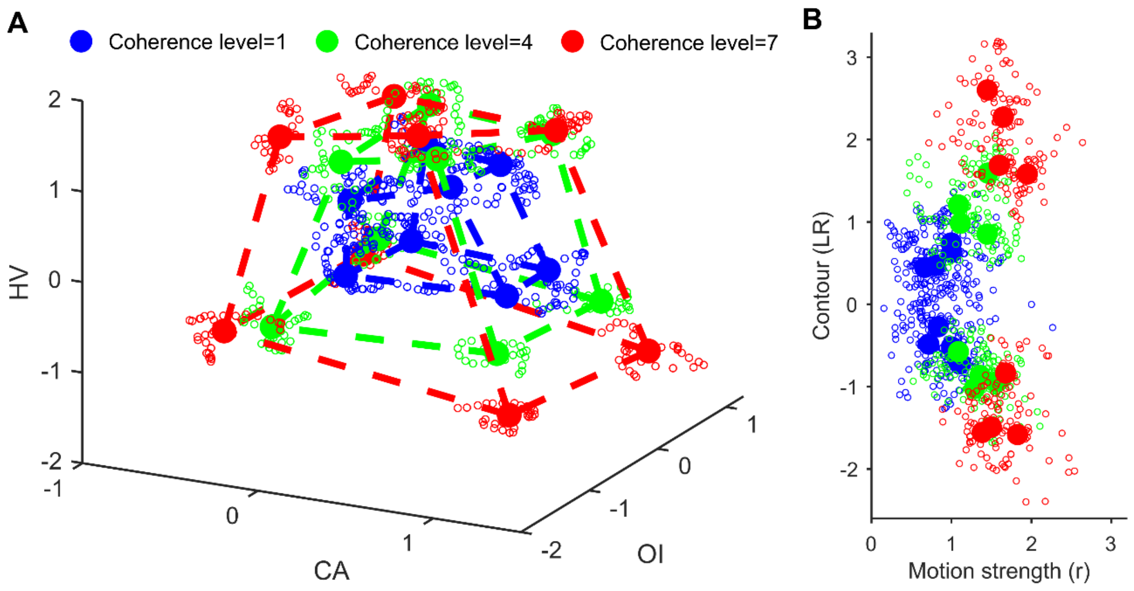


**Supplementary Figure 8. Neural geometries for different coherence levels.**

(A) Sensory manifolds in a 3-D space constructed by the three stimulus dimensions for different coherence levels. Vertices on sensory manifolds elicited by different coherence levels showed the geometry of concentric cubes when clusters' centroids (filled circles) at each coherence level were connected by dotted lines. Colors represent different coherence levels. The results for each monkey were shown in Supplementary Fig. 9 C&D. (B) The projection of neural data onto the LR axis. The vertical axis represents the projections of the neural states to the LR axis. The horizontal axis represents the distance ($r$) between the origin and the neural states shown in panel A, reflecting the magnitude of the neural population in response to the coherence level. Note that the horizontal axis shown here did not imply its existence in the neural space; it was presented here for illustrative purpose only.

### N.9. Results for individual monkeys

The main findings were also present in each individual monkey.

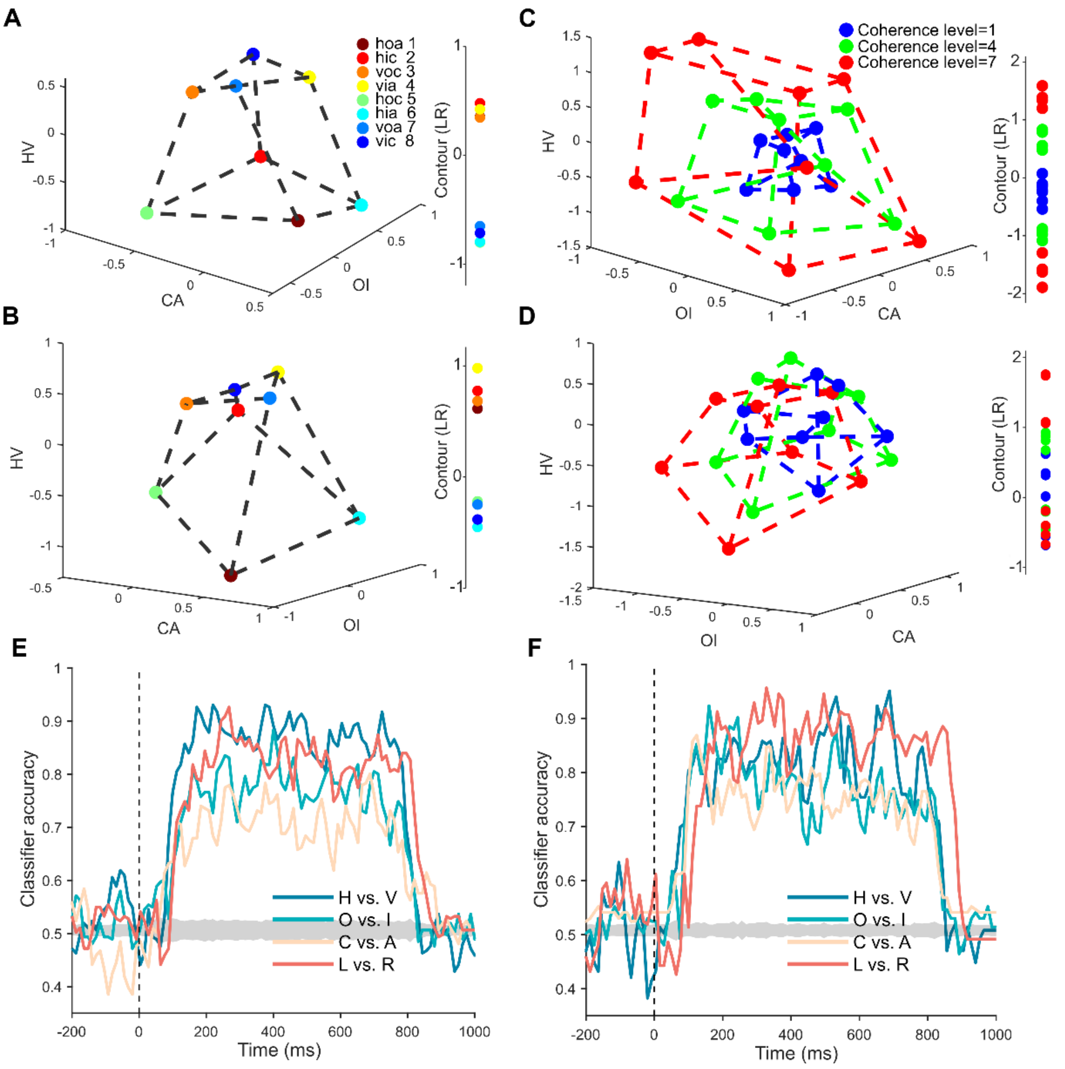


**Supplementary Figure 9. Sensory manifolds and SVM results for each individual monkey.**

(A-B) The projection of neural states in the steady phase into the 3-D subspace constructed by the three stimulus dimensions and the 1-D perceptual dimension (i.e., MIC orientations). Only the centroids of the neural states elicited by the 8 stimuli were shown. Panel A and B show the results for Monkey S and W, respectively.

(C-D) The effect of motion coherence on sensory and perceptual manifolds, respectively. Panel C and D show the results for Monkey S and W, respectively.

(E-F) The time course of SVM classifiers' overall accuracies. Panel E and F show the results for Monkey S and W, respectively.

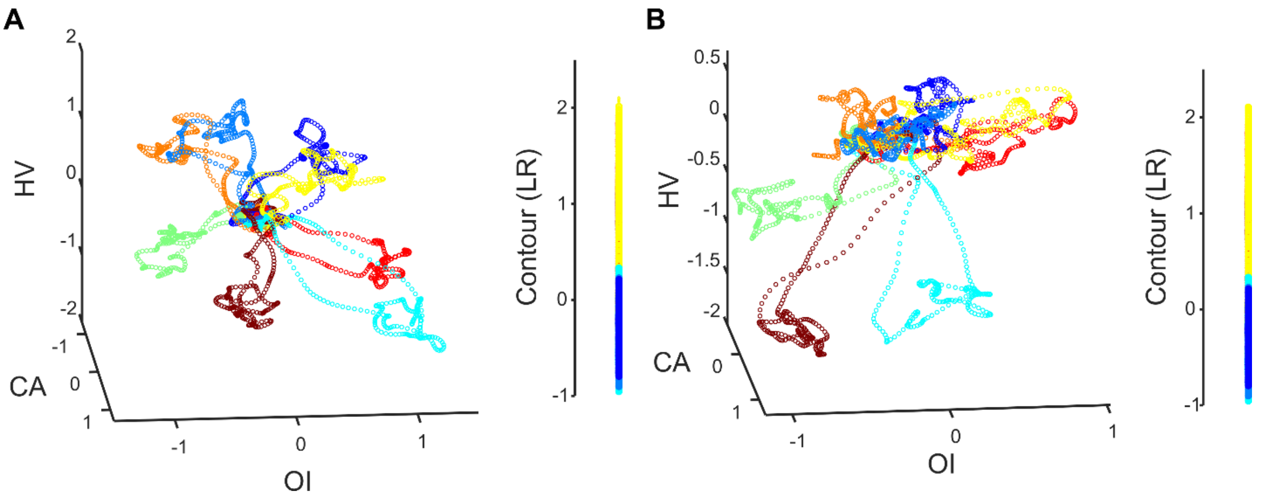


**Supplementary Figure 10. Neural trajectory at coherence level 7 for each individual monkey.** Panel A and B represent the neural trajectories at a coherence level of 7 for Monkey S and Monkey W, respectively. In each panel, the 3-D subspace (left) depicts the stimulus neural trajectory, and the 1-D subspace (right) depicts the projection of the neural trajectory onto the contour dimension.

### N.10. Temporal response of neural manifold

To better illustrate the dynamic properties of the neural manifolds, we show the time courses of the neural states projected to the sensory and perceptual axes (Supplementary Fig. 11A–D). Furthermore, we demonstrate snapshots of the dynamic neural manifold (Supplementary Fig. 11E–I).

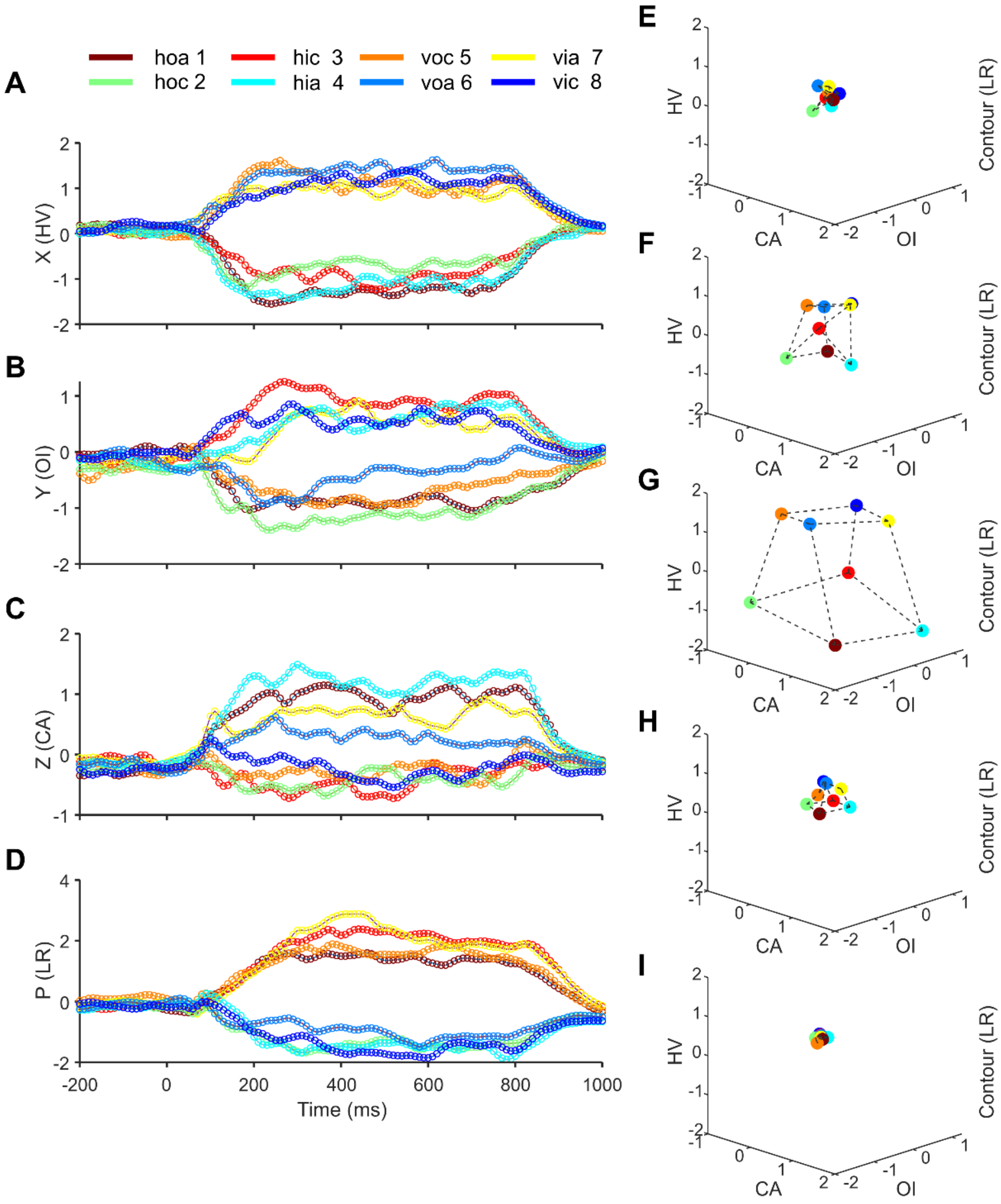


**Supplementary Figure 11. Neural geometry dynamics in embedded space.**

(A–D) The time courses of neural states (-200-1000ms) projected to the three sensory axes and one perceptual axis. Colors represent 8 different stimuli. These axes were selective to the corresponding dimension information.

(E–I) In the 3-D subspace constructed by the three stimulus dimensions (left) and the 1-D subspace by LR axis (right), the neural manifold at different time points (80, 120, 300, 900, 1000ms, respectively) is shown. Data with a coherence level 7 were used here. After stimulus onset, the manifold expanded gradually from the vicinity of the origin (panel E) to a cubic shape (panel G). Then, after stimulus offset, the cubic shape collapsed back to the origin (panel I).

### N.11. Additional main axes

**The double-twisting model.**

In Fig. 4C&D, the double-twist model is illustrated using the $v$ axis, which is the 1-twist axis. Here, we show other 1-twist axes, the $u$ and $w$ axes, and their existence in the neural space in Supplementary Fig. 12.

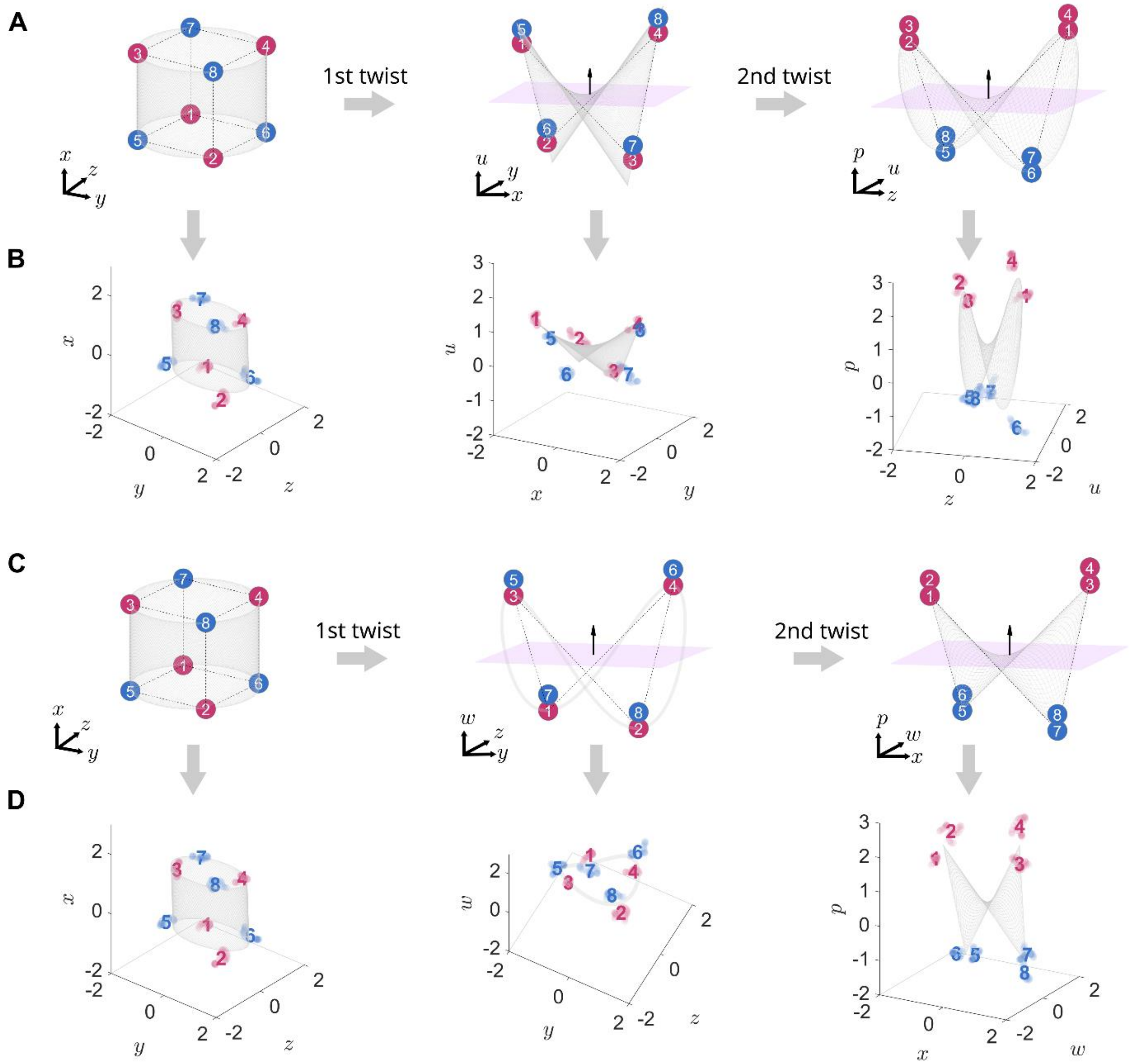


**Supplementary Figure 12. Two other routes in the double-twisting model.**

(A) One-twist axis $u$, equivalent to the logical operation $U = X \oplus Y$ (middle); two-twist axis $p$, equivalent to the logical operation $P = U \oplus Z$ (right).

(B) The neural data were projected to the corresponding subspaces and were fitted to the manifolds presumed by the double-twist operations. The existence of the $u$ axis is evident in the middle panel.

(C) One-twist axis $w$, equivalent to the logical operation $W = Y \oplus Z$ (middle); two-twist axis $p$, equivalent to the logical operation $P = W \oplus X$ (right).

(D) Same as panel B except the data are for the route in panel C. The existence of the $w$ axis is evident in the middle panel.

**Comparison between the empirical and theoretical $p$ axes.**

When the manifolds derived from double-twist operation was fitted to the neural data, its theoretical $p$ axis was adjusted by the transformation used in the fitting. It therefore provided an opportunity to compare the adjusted theoretical $p$ axis with the empirical $p$ axis obtained directly from applying SVM to the neural data. The empirical and theoretical $p$ axes were expected to be close to each other, and they are as shown in Supplementary Fig. 13.

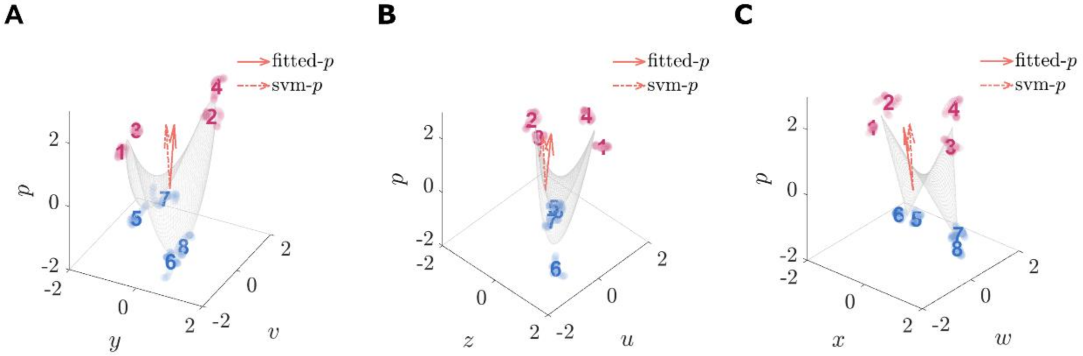


**Supplementary Figure 13. Comparison between the empirical $p$ axis from the neural data and the theoretical $p$ axis from the double-twist operations.** (A)-(C) Three 3-D subspaces incorporating the empirical $p$ axis (i.e., the SVM-$p$ axis) and two other axes were constructed for illustration. By fitting the manifold obtained from double-twist operations to the neural states within these 3-D subspaces using the least-square method, a linear transformation matrix $F$ was generated with one of its column vectors representing the theoretical $p$ axis (i.e., the fitted-$p$ axis). Evidently, in these subspaces, the theoretical $p$ axis was approximately parallel to the empirical $p$ axis. To further quantify this observation, angles between the theoretical and empirical $p$ axes were calculated across all 35 possible 3-D subspaces specified by the double-twist operations, and the angles between these empirical $p$ axes and the theoretical $p$ axis were consistently small (mean = 6.5°, std = 1.8°).

**Properties of the main axes.**

Linear SVMs were used to find the 7 main axes: the $x$, $y$, $z$, $u$, $v$, $w$, and $p$ axes. Properties including classification accuracies, classification directions, and classification dynamics, were reported by SVMs. Supplementary Fig. 14 shows these properties for the 7 main axes. Classification accuracies shown in Fig. 2A are not reported for succinctness.

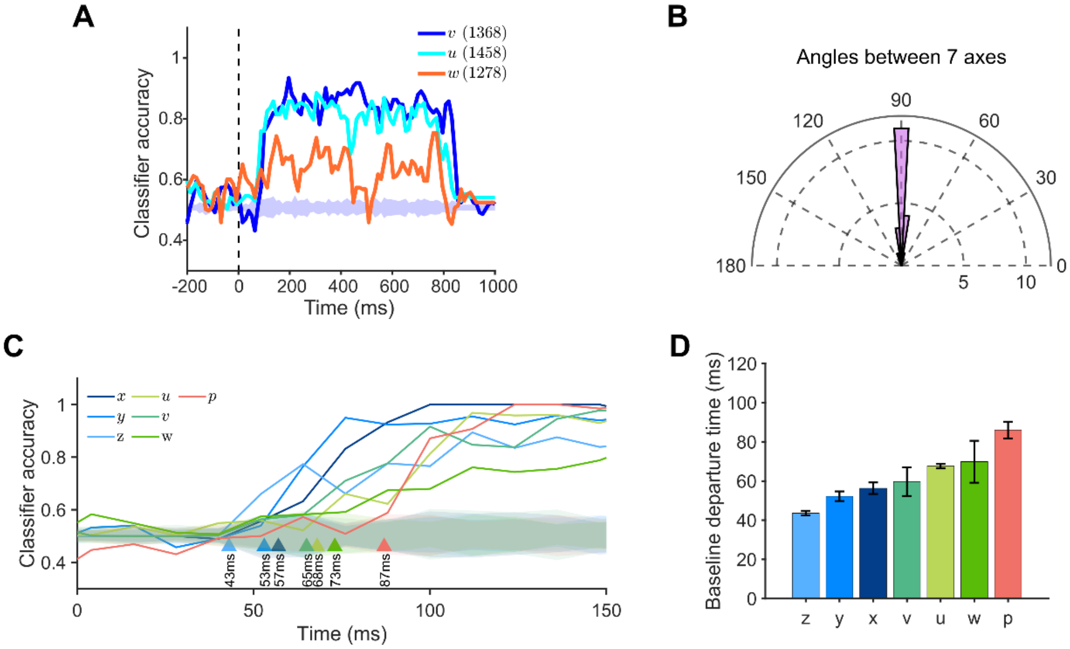


**Supplementary Figure 14. The accuracies, angles, and emergence timing of the main axes.**

(A) The linear SVM classification accuracies of the intermediate axes: $u$, $v$, $w$. The corresponding four stimuli, e.g., {1, 3, 6, 8}, that belonged to one class were shown in the parentheses. The shaded area shows the baseline accuracy generated by randomly shuffling the class labels. Note that despite the classification accuracy of the $w$ axis being lower than other axes' accuracies, it was higher than the baseline for most of stimulus presentation time.

(B) The distribution of the angles between any pair of the 7 main axes, including the $x$, $y$, $z$ and $p$ axes in Fig. 2A and $u$, $v$, and $w$ in panel A. All the angles concentrated around 90°. Thus, the 7 main axes were mutually orthogonal.

(C) To examine whether the emerging latencies differed significantly across axes, we randomly divided the neural data into training and testing sets 100 times to obtain SVM classification accuracies for each axis. The emerging latency of an axis was defined as the time point where its classification accuracy exceeded the shuffled baseline and remained above it during stimulus presence. Evidently, the latencies for the $z$, $y$, and $x$ axes were the shortest (no twist occurred); the $v$, $u$, and $w$ axes had intermediate latencies (after 1 twist); and the p axis had the longest latency (after 2 twists).

(D) Pairwise t-tests (Bonferroni corrected) showed that the latency differences between any two axes adjacent in the ranking order were significant ($ps < 0.001$), except for the $u$ and $w$ axis difference ($p = 0.08$). Therefore, the ranking order of the emergence among individual axes aligned with the latency ranking order among the 0-twist, 1-twist, and 2-twist axes reported in the study.

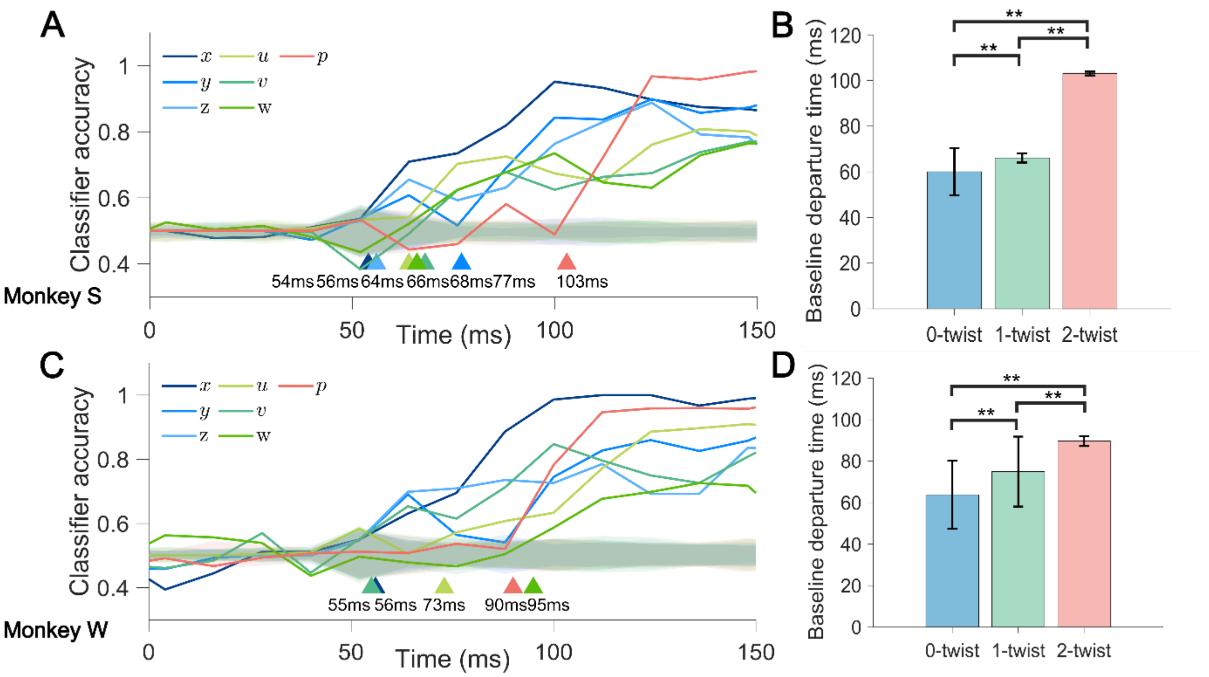


**Supplementary Figure 15. The emergence timing of the main axes for single monkey.**

(A-D) Similar to Supplementary Figure 14 but for single monkey. A and C: Time courses of classification accuracies of the axes for single monkey. Shaded regions denote the mean SVM accuracy (±3 standard deviations) of the baselines by shuffling the class labels (200 times for each classification). B and D: The latencies of the 0-twist axes (blue), the 1-twist axes (green), and the 2-twist axis (red). **: $p<0.001$. A and B are Monkey S results, C and D are Monkey W results. Due to the limited number of neurons per monkey and noisy signal, the results for individual axes are not very clear. However, the overall trend support that the 0-twist axes were the shortest, followed by the 1-twist axes, and finally the 2-twist axis.

### N.12. The general property of V2: seven main axes in different V2 stripes.

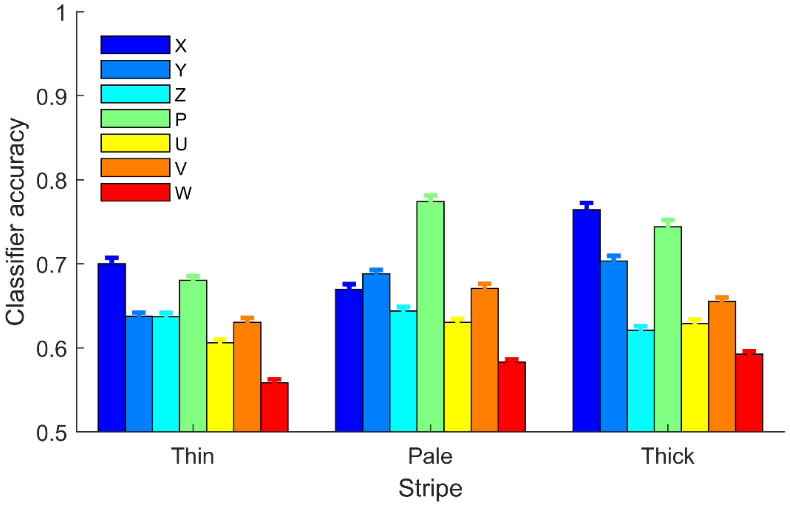


**Supplementary Figure 16. The representation of the 7 axes in thin, pale, and thick stripes of the V2 area, respectively.** In the V2 area, there are three types of cytochrome oxidase (CO) stripes – thin, thick, and pale – each associated with relatively distinct visual processing functions like color, motion and depth perception, spatial orientation and form, respectively. The 93 neurons were divided into three groups according to their respective stripe types: thin (n = 19), thick (n = 39), and pale (n = 35), based on the recording sites in the V2 area (Ma et al., 2021). SVM analyses identical to those in the study were performed to assess classification accuracies for the 7 axes (each repeated 100 times) in the three stripes. While the classification accuracies were generally lower due to the smaller number of neurons for each stripe, the 7 axes were identified in each of three CO stripes ($p$s < 0.001), with similar accuracy distributions ($r$s > 0.7, $p$s < 0.05). Therefore, the findings in the study likely reflect a general property of V2 neurons.

### N.13. Linearly separable classifications

**The number of linearly separable classifications.**

The number of linearly separable classifications is 104. We obtained this number by enumeration. Supplementary Fig. 17 shows how the number of linear separability was enumerated.

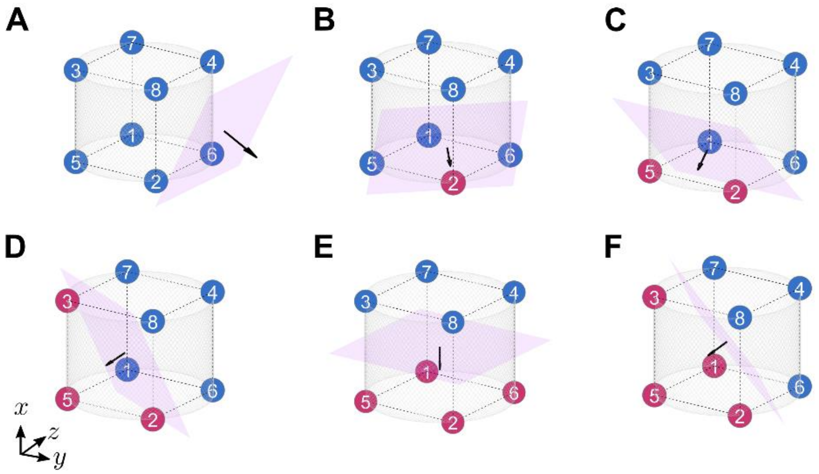


**Supplementary Figure 17. The number of linear binary classifications in the 3-D stimulus space.**
A classification is denoted by subsets $X$ versus $X^c$ , and its symmetric classification is $X^c$ versus $X$. We only consider one side of the symmetry.
(A) When $\|X\| = 0$, there is 1 classification.
(B) When $\|X\| = 1$, there are 8 classifications for isolating each vertex.
(C) When $\|X\| = 2$, there are 12 classifications for isolating each edge.
(D) When $\|X\| = 3$, there are 24 classifications for 4 different ways of isolating 3 vertices on 6 faces.
(E) When $\|X\| = 4$, in this parallel case, there are 3 classifications for isolating a complete face.
(F) When $\|X\| = 4$, in this corner case, there are 4 classifications for isolating a complete corner.
Overall, there are $(1 + 8 + 12 + 24 + 3 + 4) \times 2 = 104$ different linear binary classifications.

**Enumeration of linearly separable classifications.**

In Supplementary Table. 1, we listed all possible linearly separable classifications. Since the classifications are symmetric, there are 52 unique classifications listed in the stimulus space. The number of classifications was enumerated as in Supplementary Fig. 17. The SVM analysis on the synthesized pure selectivity neural data (Supplementary Fig. 21) confirmed that the classifications listed in this table are indeed separable, with a minimal accuracy of 95.7% when the box constraint was set at 1.

**Supplementary Table 1. The 104 linearly separable classifications in the stimulus space.**

| Subspace $y$-$z$-$x$ | | | | | |
|---|---|---|---|---|---|
| ∅ vs. | 15 vs. 234678 | 46 vs. 123578 | 156 vs. 23478 | 268 vs. 13457 | 1235 vs. 4678 |
| 12345678 | 16 vs. 234578 | 47 vs. 123568 | 157 vs. 23468 | 347 vs. 12568 | 1246 vs. 3578 |
| 1 vs. 2345678 | 17 vs. 234578 | 48 vs. 123567 | 167 vs. 23458 | 348 vs. 12567 | 1256 vs. 3478 |
| 2 vs. 1345678 | 25 vs. 134678 | 125 vs. 34678 | 235 vs. 14678 | 357 vs. 12468 | 1347 vs. 2568 |
| 3 vs. 1245678 | 26 vs. 134578 | 126 vs. 34578 | 238 vs. 14567 | 358 vs. 12467 | 1357 vs. 2468 |
| 4 vs. 1235678 | 28 vs. 134567 | 135 vs. 24678 | 246 vs. 13578 | 378 vs. 12456 | 1467 vs. 2358 |
| 5 vs. 1234678 | 35 vs. 123678 | 137 vs. 24568 | 248 vs. 13567 | 467 vs. 12358 | 1567 vs. 2348 |
| 6 vs. 1234578 | 37 vs. 124568 | 146 vs. 23578 | 256 vs. 13478 | 468 vs. 12357 | |
| 7 vs. 1234568 | 38 vs. 124567 | 147 vs. 23568 | 258 vs. 13467 | 478 vs. 12356 | |
| 8 vs. 1234567 | | | | | |

### N.14. Linearly inseparable classifications

**Subspaces of the theoretically-derived perceptual manifold.**

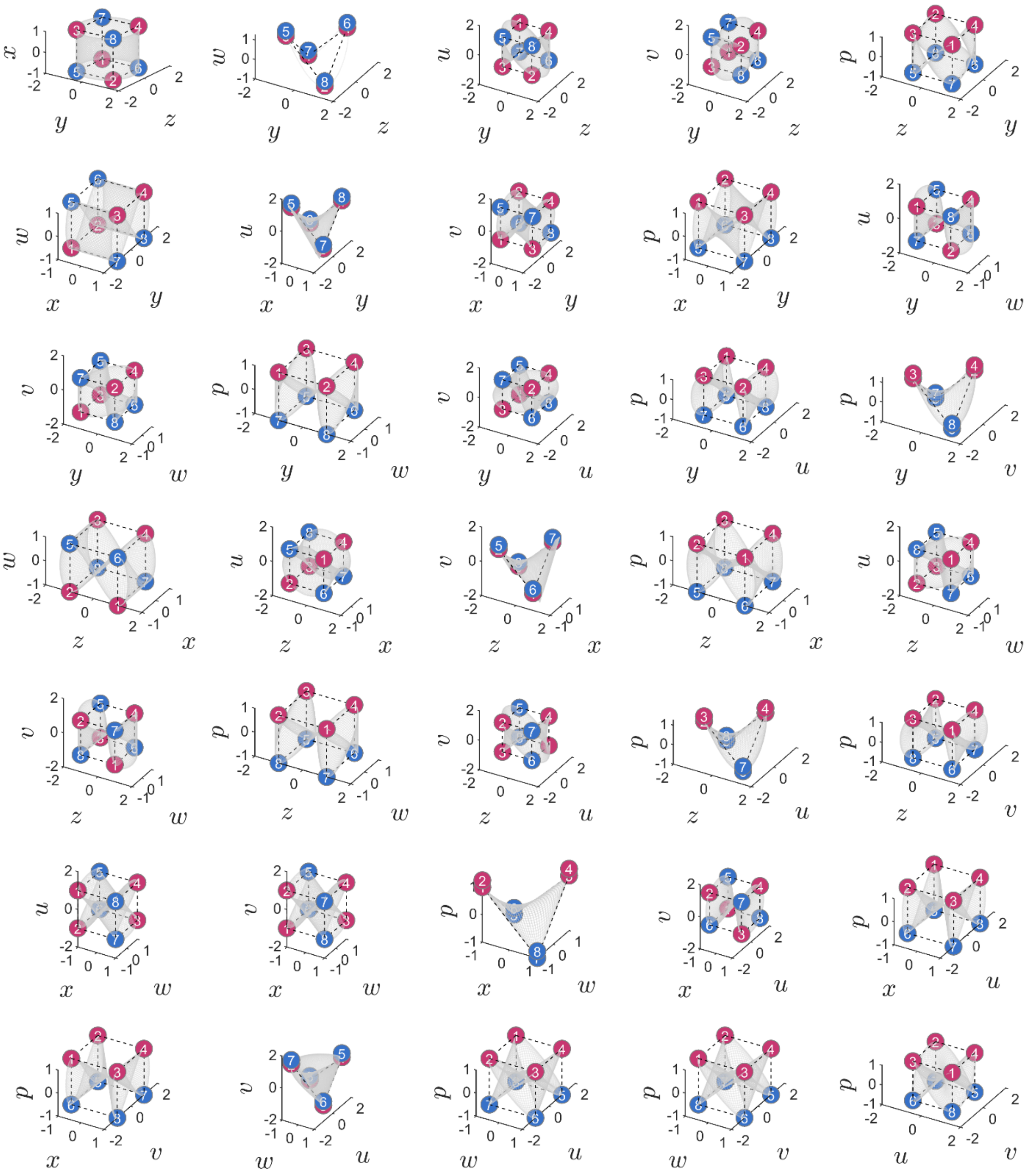

**Supplementary Figure 18. A complete view of the theoretical perceptual manifold.**
In the 7-D space generated by the double-twist operations, all the linearly inseparable classifications in the 3-D stimulus space becomes linearly separable. By selecting 3 axes of the 7-D space, in total $\binom{7}{3} = 35$ different 3-D subspaces can be constructed. Projections of the perceptual manifold are shown in these subspaces. Each linearly inseparable classification of the stimuli in the stimulus space can find its linearly separable version in one of the 35 subspaces (Supplementary Table. 2). Together, the projections in the 35 subspaces give a complete view of the theoretical perceptual manifold. Stimuli in the red vertices show the right-tilted contour, and those in the blue vertices show the left-titled contour.

**Enumeration of linearly inseparable classifications.**

In Supplementary Table. 2, we list 9 subspaces that have already provided 152 linearly separable versions of the classifications that are linearly inseparable in the stimulus space. Since many classifications are symmetric, we list 152/2=76 unique classifications in 9 subspaces after excluding the symmetric ones. The subspaces are shown in Supplementary Fig. 18. The 9 subspaces all require at least one of the $u$, $v$, $w$, and $p$ axes. Linear classifications in other subspaces are replicates of the ones listed here.

**Supplementary Table 2. The 152 linearly inseparable classifications become linearly separable in new subspaces obtained with the double-twist model.**

| Subspace $y$-$z$-$p$ | | Subspace $y$-$z$-$u$ | | Subspace $y$-$z$-$v$ | |
|---|---|---|---|---|---|
| 12 vs. 345678 | 234 vs. 15678 | 14 vs. 235678 | 237 vs. 14568 | 57 vs. 123468 | 247 vs. 13568 |
| 13 vs. 245678 | 567 vs. 12348 | 58 vs. 123467 | 267 vs. 13458 | 68 vs. 123457 | 257 vs. 13468 |
| 23 vs. 145678 | 568 vs. 12347 | 67 vs. 123458 | 367 vs. 12458 | 136 vs. 24578 | 368 vs. 12457 |
| 24 vs. 135678 | 578 vs. 12346 | 145 vs. 23678 | 458 vs. 12367 | 138 vs. 24567 | 457 vs. 12368 |
| 123 vs. 45678 | 678 vs. 12345 | 148 vs. 23567 | 1358 vs. 2467 | 168 vs. 23457 | 1457 vs. 2368 |
| 124 vs. 35678 | 1234 vs. 5678 | 158 vs. 23467 | 1367 vs. 2458 | 245 vs. 13678 | 1468 vs. 2357 |
| 134 vs. 25678 | | 236 vs. 14578 | | | |
| **Subspace $y$-$w$-$p$** | | **Subspace $z$-$x$-$w$** | | **Subspace $z$-$w$-$v$** | |
| 1237 vs. 4568 | | 34 vs. 125678 | 345 vs. 12678 | 18 vs. 234567 | 1238 vs. 4567 |
| 1248 vs. 3567 | | 56 vs. 123478 | 346 vs. 12578 | 27 vs. 134568 | 1247 vs. 3568 |
| 1345 vs. 2678 | | 78 vs. 123456 | 356 vs. 12478 | 36 vs. 124578 | 1346 vs. 2578 |
| 1578 vs. 2346 | | 127 vs. 34568 | 1258 vs. 3467 | 45 vs. 123678 | 1678 vs. 2345 |
| | | 128 vs. 34567 | 1267 vs. 3458 | | |
| | | 178 vs. 23456 | 1456 vs. 2378 | | |
| | | 278 vs. 13456 | 1478 vs. 2356 | | |
| **Subspace $x$-$y$-$w$** | | **Subspace $x$-$u$-$p$** | | **Subspace $w$-$u$-$v$** | |
| 456 vs. 12378 | | 1236 vs. 4578 | | 1278 vs. 3456 | |
| 1257 vs. 3468 | | 1245 vs. 3678 | | 1368 vs. 2457 | |
| 1268 vs. 3457 | | 1348 vs. 2567 | | 1458 vs. 2367 | |
| 1356 vs. 2478 | | 1568 vs. 2357 | | | |
| 1378 vs. 2456 | | | | | |

**Neural data visualization in perceptual subspaces.**

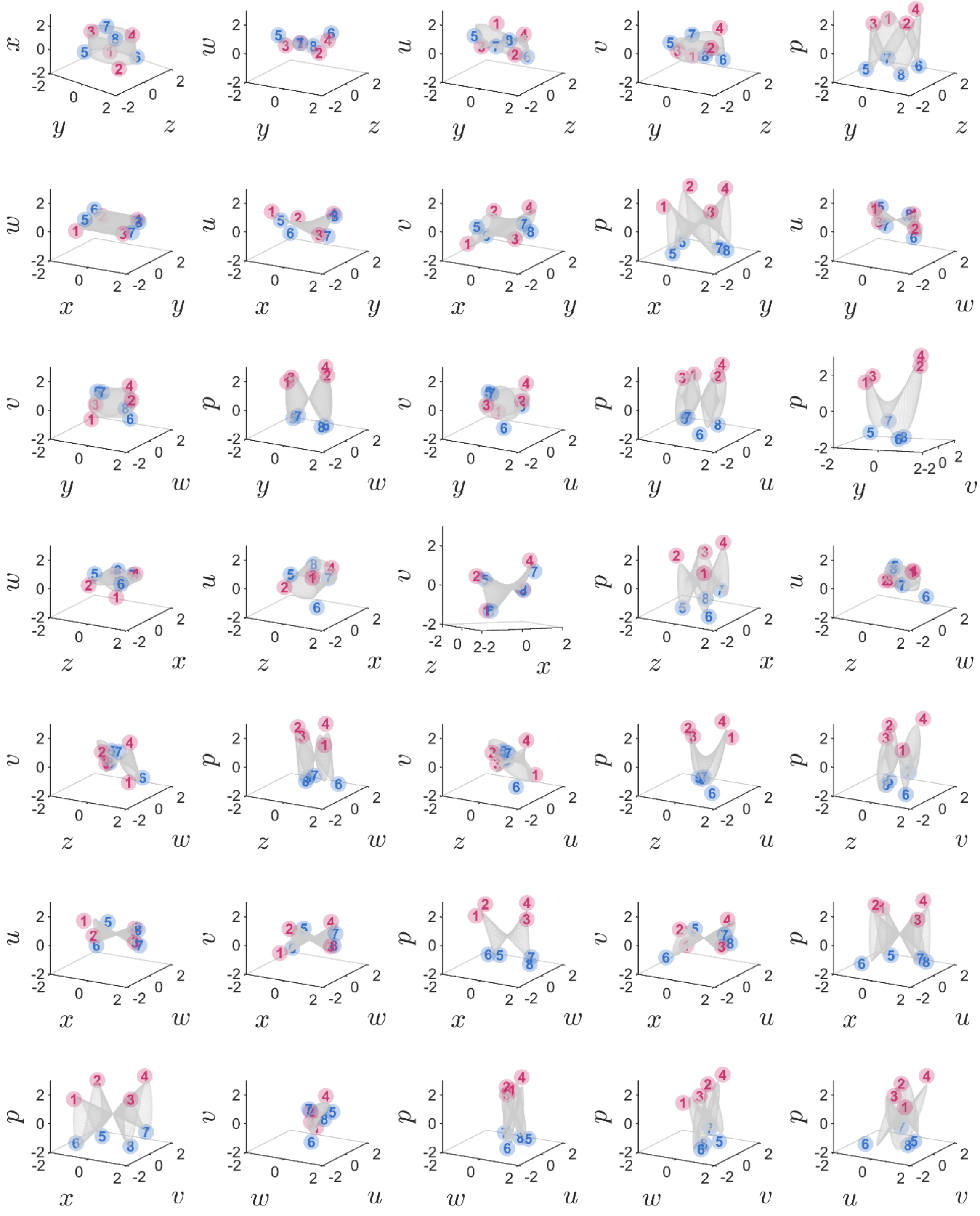

**Supplementary Figure 19. The neural data in 35 different 3-D perceptual subspaces.**

. Corresponding to the 35 different 3-D subspaces generated by the double-twist operations, we can visualize the neural data in the 35 perceptual subspaces. The subspaces for visualization were created with the 7 main axes found by linear SVMs, because these axes were mutually orthogonal (Supplementary Fig. 14B). For subspaces created by axes with high classification accuracies, the consistency between the data and the theoretically derived perceptual manifold was high, for instance, in the $y$-$z$-$p$ and $y$-$v$-$p$ subspaces. For subspaces created by axes with low classification accuracies, the consistency was relatively low. For instance, because the separation margin of the $w$ axis was short, the data-model consistency in the $x$-$y$-$w$ and $z$-$w$-$u$ spaces was relatively small, suggesting that the dimension of the $w$ axis might be less important in the perceptual manifold. Nevertheless, we still can visualize the linear separation in these perceptual subspaces. For clarity, only centers of the point clusters are shown. The red vertices have a right-tilted contour (R), and the blue ones have a left-tilted contour (L).

### N.15. Independence of classifications

**Definition of independent classification.**

Denote a classification as grouping a set of data points into two subsets: a subset $X$ versus the complement $X^c$. Two independent classifications of the same set, $X$ versus $X^c$ and $Y$ versus $Y^c$, have the property that knowing an object $a$ belonging to $X$ does not affect the probability of correctly guessing whether $a$ belongs to $Y$. That is, the two classifications are completely uncorrelated.

The classifications for HV, OI, and CA are indeed mutually independent (see below). Their classification directions in the neural space were mutually orthogonal. Intriguingly, the orthogonality corresponded nicely to the independence of classifications. This correspondence motivated us to define mathematically what dependent and independent classifications are.

Here, we explain the conditions for two different classifications to be independent. Suppose the set $U$ contains all the data points. Non-empty and non-overlapping subsets cover $U$: $A \cup B \cup C \cup D = U$. Let the first binary classification be $X = A \cup B$ versus $X^c = C \cup D$. Let the second binary classification be $Y = A \cup C$ versus $Y^c = B \cup D$. We then get $X \cap Y = A$, $X \cap Y^c = B$, $X^c \cap Y = C$, and $X^c \cap Y^c = D$. For an arbitrary data point $a$, we derive the condition under which $P(a \in Y | a \in X) = P(a \in Y)$. According to Bayes' rule, we derive

$$P(a \in Y | a \in X) = \frac{P(a \in Y, a \in X)}{P(a \in X)} = \frac{||X \cap Y||}{||X||} = \frac{||A||}{||A \cup B||} = \frac{||A||}{||A|| + ||B||}.$$

On the other side, we derive

$$P(a \in Y) = \frac{||Y||}{||U||} = \frac{||A \cup C||}{||A \cup B \cup C \cup D||} = \frac{||A|| + ||C||}{||A|| + ||B|| + ||C|| + ||D||}.$$

Equalizing these two expressions and simplifying, we obtain

$$\frac{||A||}{||B||} = \frac{||C||}{||D||}.$$

Furthermore, if we additionally have $||X|| = ||X^c||$ and $||X|| = ||Y||$, we can derive $||A|| = ||B|| = ||C|| = ||D||$. In our work, given one classification $X$ and $X^c$ with $||X|| = ||X^c|| = 4$, if we swap 2 elements in $X$ with 2 elements in $X^c$, we get a new classification $Y$ and $Y^c$ that is independent to the original one.

**Properties of dependent and independent classifications.**

With respect to binary classifications where each class has 4 elements, there are $\binom{8}{4}/2 = 35$ different classifications, after excluding symmetric ones. As discussed above, any two classifications are either independent or dependent. Using linear SVMs, we computed the accuracies of these classifications and found classification directions of these separations. All the classification directions existed according to the classification accuracies (Supplementary Fig. 20A). We computed the angles between the classification directions of the independent classifications and the angles between the classification directions of the dependent classifications and showed their distribution in Supplementary Fig. 20B.

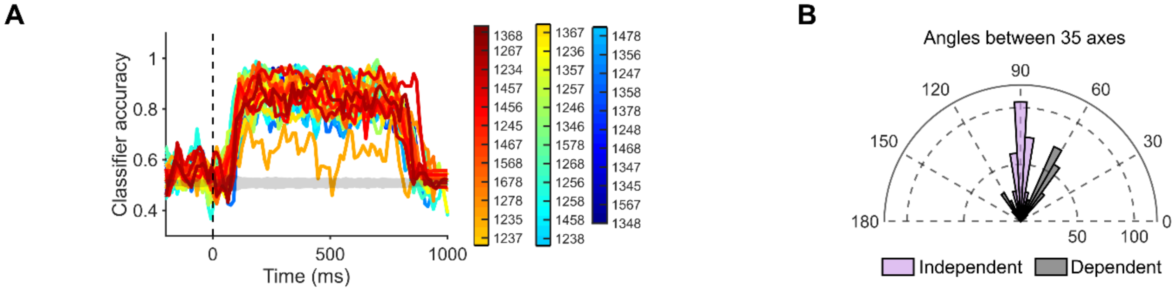


**Supplementary Figure 20. Existence of dependent and independent classification directions and the angle distribution.**

(A) The 35 classification accuracies of all 4-element binary classifications. The trajectories are color coded and labeled by the four stimulus conditions in one class. All the classification accuracies are above the baseline (shaded area), except the classification 1278, which is the $w$ axis, was slightly worse as in Supplementary Fig. 14A.

(B) The distribution of the angles between any pair of the 35 classification directions. The angles between classification directions of any two independent classifications are shown in purple, whereas the angles between classification directions of any two dependent classifications are shown in gray. The angles of independent classification directions concentrated around 90°, while the angles of dependent classification directions concentrated around either 60° or 120°.

### N.16. Testing nonlinear mixed selectivity and synthesizing purely selective neurons

**ANOVA as a tool for testing nonlinear mixed selectivity.**

When a neuron has *linear* mixed selectivity to any two features, including linear pure selectivity as its special case, here we show that the output of the neuron is subject to no effects of interaction between the features in ANOVA tests. Denote the output of a neuron as $s$ and two features as $a$ and $b$. Let $a$ have $i = 1, \dots, m$ levels and $b$ have $j = 1, \dots, n$ levels. Since the derivation of interaction effects only requires between-group variations, we assume every level combination of features $a$ and $b$ has one trial, thus ignoring within-group variations without losing generality.

Linear mixed selectivity means that the neuron linearly mixing $a_i$ and $b_j$ has its output being $s_{ij} = \alpha_a a_i + \alpha_b b_j$, where $\alpha_a$ and $\alpha_b$ are two constants. Define $\bar{s}$ as the grand mean, i.e., $mn\bar{s} = \sum_{ij} s_{ij}$; $\bar{s_i}$ as the mean for $a = a_i$, i.e., $n\bar{s_i} = \sum_j s_{ij}$; and $\bar{s_j}$ as the mean for $b = b_j$, i.e., $m\bar{s_j} = \sum_i s_{ij}$. From the definitions we can derive $m\bar{s} = \sum_i \bar{s_i}$ and $n\bar{s} = \sum_j \bar{s_j}$. In ANOVA, the sum of squares (SS) of interaction effects is

$$SS_{\text{interact}} = \sum_{ij} \left(s_{ij} - \bar{s}\right)^2 - \sum_{ij} (\bar{s_i} - \bar{s})^2 - \sum_{ij} \left(\bar{s_j} - \bar{s}\right)^2.$$

Our goal is to show $SS_{\text{interact}} = 0$.

To do this, we expand the above equation and use the definitions of $\bar{s}$, $\bar{s_i}$, and $\bar{s_j}$ to get

$$SS_{\text{interact}} = \sum_{ij} \left(s_{ij}^2 - 2 s_{ij}\bar{s} + \bar{s}^2\right) - \sum_{ij} \left(\bar{s_i}^2 - 2\,\bar{s_i}\bar{s} + \bar{s}^2\right) - \sum_{ij} \left(\bar{s_j}^2 - 2\,\bar{s_j}\bar{s} + \bar{s}^2\right)$$

$$= \sum_j \sum_i \left(s_{ij}^2 - \bar{s_i}^2 + \bar{s}^2 - \bar{s_j}^2\right)$$

$$= \textstyle\sum_j \left(\sum_i \left(s_{ij} - \bar{s_i}\right)\left(s_{ij} + \bar{s_i}\right) + \sum_i \left(\bar{s} - \bar{s_j}\right)\left(\bar{s} + \bar{s_j}\right)\right).$$

Since $s_{ij} = \alpha_a a_i + \alpha_b b_j$, we can derive $s_{ij} - \bar{s_i} = \alpha_a a_i + \alpha_b b_j - \alpha_a a_i + \frac{\alpha_b}{n} \sum_j b_j = \alpha_b (b_j + \frac{1}{n} \sum_j b_j)$ which is independent of $i$. Also, $\left(\bar{s} - \bar{s_j}\right)$ is independent of $i$. We can take $\left(s_{ij} - \bar{s_i}\right)$ and $\left(\bar{s} - \bar{s_j}\right)$ out of the summation over $i$. Hence,

$$SS_{\text{interact}} = \sum_j \left( \left(s_{ij} - \bar{s_i}\right) \sum_i \left(s_{ij} + \bar{s_i}\right) + \left(\bar{s} - \bar{s_j}\right) \sum_i \left(\bar{s} + \bar{s_j}\right) \right).$$

Because $\sum_i s_{ij} = m\overline{s_j} = \sum_i \overline{s_j}$ and $\sum_i \overline{s_i} = m\bar{s} = \sum_i \bar{s}$, we get $\sum_i (s_{ij} + \overline{s_i}) = \sum_i (\bar{s} + \overline{s_j})$. Hence,

$$SS_{\text{interact}} = \sum_j \left( (s_{ij} - \overline{s_i} + \bar{s} - \overline{s_j}) \sum_i (\bar{s} + \overline{s_j}) \right).$$

Since $(s_{ij} - \overline{s_i})$ and $(\bar{s} - \overline{s_j})$ are independent of $i$ as proved above, their sum $(s_{ij} - \overline{s_i} + \bar{s} - \overline{s_j})$ is independent of $i$. We can move it back into the summation over $i$. Also, we see $(\bar{s} + \overline{s_j})$ is independent of $i$, we can take it out of the summation over $i$. Therefore, we arrive at

$$SS_{\text{interact}} = \sum_j (\bar{s} + \overline{s_j}) \left( \sum_i s_{ij} - \sum_i \overline{s_i} + \sum_i \bar{s} - \sum_i \overline{s_j} \right).$$

Because $\sum_i \overline{s_j} = m\overline{s_j} = \sum_i s_{ij}$ and $\sum_i \bar{s} = m\bar{s} = \sum_i \overline{s_i}$, we finally get $SS_{\text{interact}} = 0$. Therefore, for any two features, if the interaction effect is nonzero, the neuron output is neither pure nor linearly mixed responses; accordingly, the output must be a nonlinearly mixed response.

**Synthesizing purely selective neurons from recorded nonlinearly mixed selective neurons.**

Using ANOVA tests, we found all the recorded neurons had nonlinearly mixed selectivity. In order to compare classification capabilities, we created purely selective neurons. (Purely selective neurons are special cases of linearly mixed selective neurons, i.e., zero interaction effects, thus in essence they are the same.) We did this by synthesizing pure neuronal responses from the raw neural data to maximally preserve data variability (Rigotti et al., 2013). Supplementary Fig. 21 shows the procedure of synthesizing one neuron purely selective to HV from a recorded example neuron.

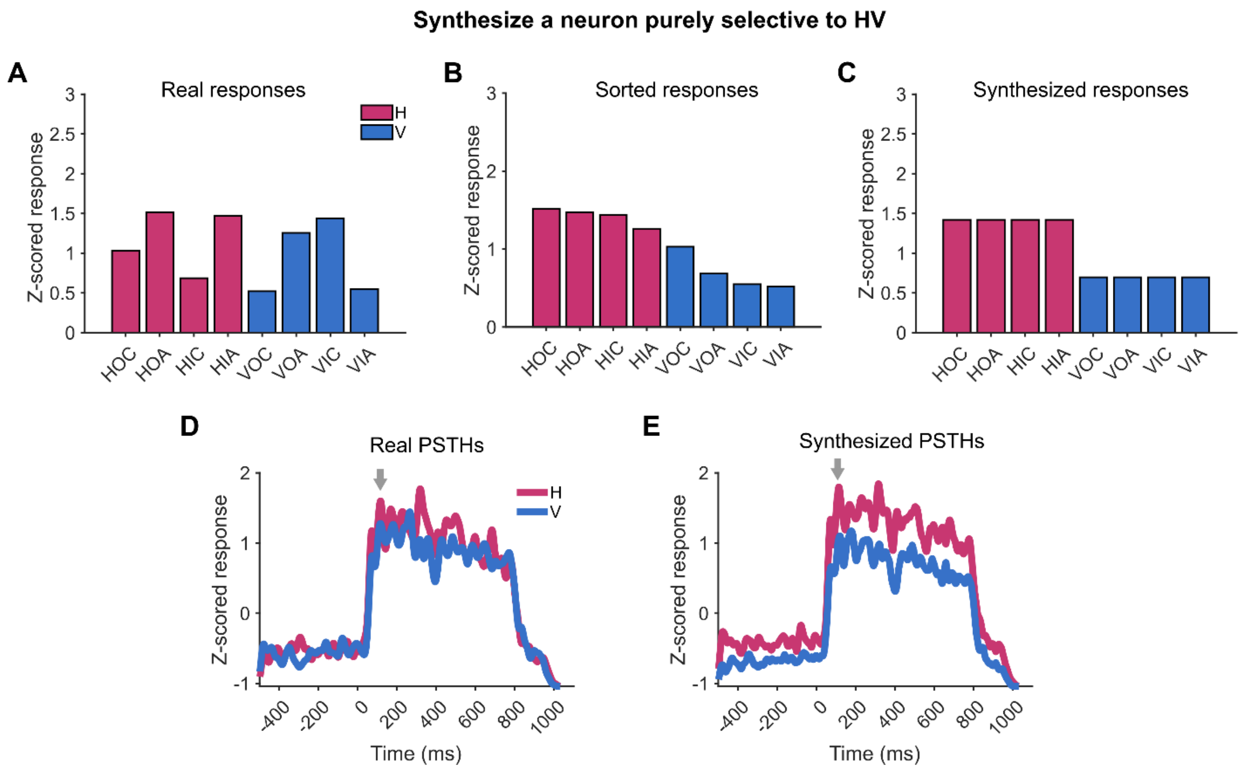


**Supplementary Figure 21. The procedure of synthesizing neurons with pure selectivity.**

(A) The activities of each neuron at each time bin were recorded for all the stimuli. The stimuli are labelled by abbreviations, e.g., HOC indicates the horizontal-outward-clockwise stimulus. Here, an example neuron's responses to different stimuli are shown. The responses are colored according to the HV dimension. The purpose was to synthesize an artificial neuron that was purely selective to HV but retaining the variability of the neuron's real responses. The process took two steps: sorting and averaging at each time bin.

(B) Step 1: The activities of all stimulus conditions were sorted regardless of the class labels. Then, the first half of the stimulus conditions were relabeled H while the second half were relabeled V.

(C) Step 2: The mean activities of each class were reassigned to every stimulus in their respective class.

(D) The actual averaged Peri-Stimulus Time Histograms (PSTHs) of the neuron in panel A. This neuron did not show evident selectivity for any stimulus dimension.

(E) The averaged PSTHs of the synthesized pure selectivity neuron. The selectivity to HV became clear. Arrows indicate the time bin during which the neuron's activities are shown in panel A, B, and C.

### N.17. Continuous manifolds can be observed in simulated network data.

In the Result section, we focused on showing the neural data of the 8 stimuli and drew the manifold predicted by the double-twist operations as the background. That is, the neural data were disconnected clusters on the manifold. To further illustrate the continuity of neural manifolds suggested by the model, we used continuous stimuli as inputs to the neural network (Fig. 7D) and analyzed the output data.

Specifically, the continuous stimulus input spanned the entire sheared configuration rings (Supplementary Fig. 1B) for the vertical and horizontal motion-axes, respectively. The diversity parameter was set at $d$=1 for the neural network. Thus, the input space was two 1-D rings (Supplementary Fig. 22A, left), with 400 samples uniformly distributed on each ring. The ring-shaped neural states corresponding to the input structure was evident in Supplementary Fig. 22B (left).

We then fitted the theoretically-derived perceptual manifold to the data points corresponding to the 8 stimuli in each 3-D subspace, which labelled either in red or in blue (Supplementary Fig. 22). Each stimulus points were computed as the average of 30 data points around the solid circles in Supplementary Fig. 22A (left). Affine transformation was used for the fitting. Similar manifold fittings were observed in other subspaces. The illustration of the simulated network data and the fitted perceptual manifolds are shown in Supplementary Fig. 22.

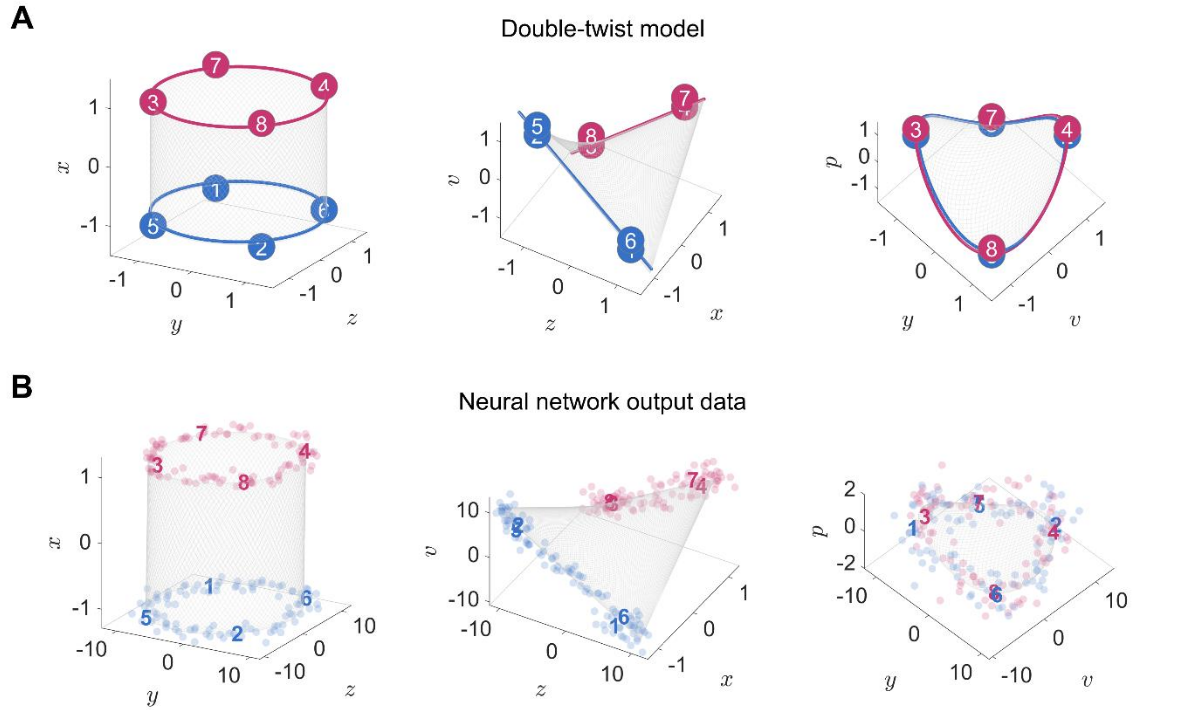


**Supplementary Figure 22. The comparison between the theoretically-derived perceptual manifold and simulated neural network data.**

(A) The theoretically-derived perceptual manifold in which the stimuli spanned the entire sheared configuration rings for the vertical and horizontal motion-axis. The continuous stimuli are highlighted in color.

Red color represents vertical motion-axis and blue represents horizontal motion-axis. Among the continuous stimuli, 8 stimuli that were tested in the experiment are marked by solid circles. Left to right represents subspaces after 0-twist, 1-twist, and 2-twist operations, respectively.

(B) The simulate network data visualized in the subspaces corresponding to panel A. The input data points were sampled from the continuous rings in the stimulus space. Left to right shows the visualization of the output data. The background manifolds were fitted to the data points corresponding to the 8 stimuli in each subspace. The rest data points positioned on the manifolds in a continuous fashion. Comparison between A and B shows close alignment of the manifold formed by the neural network to that predicted by the double-twist operations.

### N.18. Binary classification of simulated network data

**Successful binary classification for different kind classifications as a function of diversity.**

In the results presented in Fig.6D-H, we demonstrated that the number of successful classifications increased with diversity, and all binary classifications were successful when $d > 0.5$. In this context, we further separated binary classification problem into two categories: linearly inseparable and linearly separable classifications. This categorization was based on whether the classification could be linearly readout in the 3-D stimulus space (see Supplementary Fig. 17 and Supplementary Table. 1 & 2). Supplementary Fig. 23 shows that the linearly inseparable problems were concurrently resolved with the linearly separable classifications with increased amount of diversity.

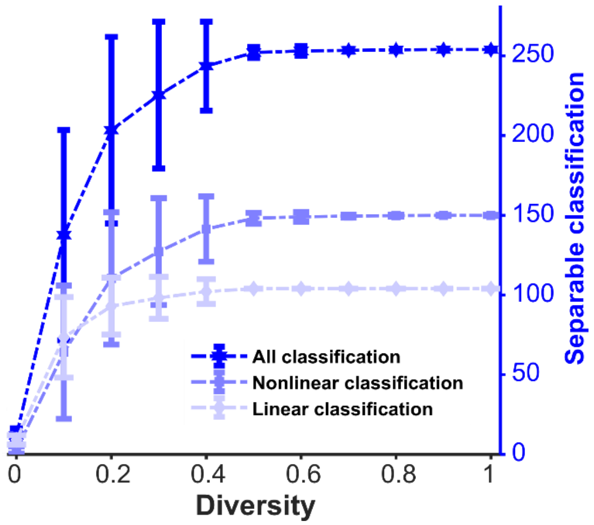


**Supplementary Figure 23. The number of successful classifications as a function of diversity.**

Three color curves represent the number of successful classifications as the function of diversity. Among these, dark blue represents all binary classification problems (256 in total), blue represents linearly inseparable binary classification problems in the 3-D stimulus space (152 in total), and light bule represents linearly separable binary classification problems in the 3-D stimulus space (104 in total).

**Successful binary classification for different kinds of neurons as a function of population size.** Supplementary Fig. 24 shows NMS neuron and pure neuron results in simulation network. Similar with Fig. 6C, pure neuron through the setting of connection weights could only solve linearly separable problems.

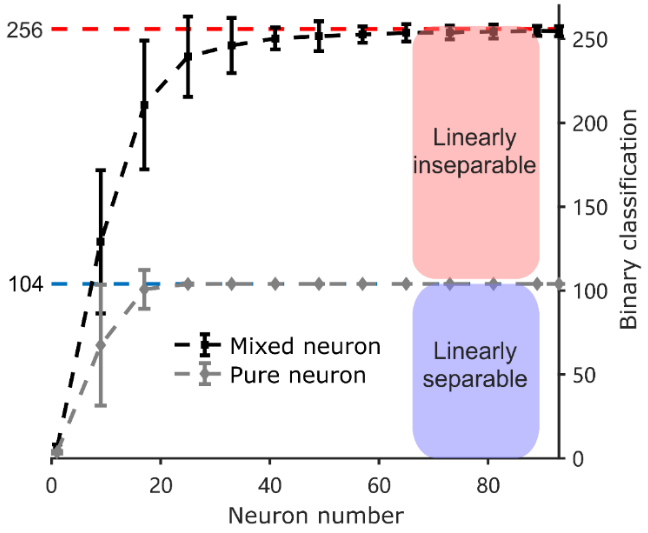


**Supplementary Figure 24. The number of successful linear classifications increases as a function of neuron population size for both pure selectivity neurons and NMS neurons.** In the neural network, the parameter $d$, representing the diversity in the connection patterns, was set to 0.6. Pure selectivity neurons were defined by their responsiveness solely to a single feature. In creating a 2-layer network similar to that shown in Fig. 6D, yet featuring pure selectivity neurons in the second layer, we adopted a similar approach to specifying connection weights as detailed in the "Connectivity patterns of neural network" section of the Methods, with a critical modification that each neuron's response was determined exclusively by a single feature to ensure pure selectivity. Simulations showed that the network with pure selectivity neurons successfully classified all linearly separable problems (104 instances in total, gray curve) but failed to solve any linearly inseparable problem. In contrast, with the same parameter $d$ (i.e., 0.6), the network with NMS neurons successfully classified both linearly separable and inseparable problems (256 instances in total, dark curve), underscoring the essential role of NMS neurons in tackling such complex problems. Error bar: standard deviation.

### N.19. Movies for stimuli and dynamic neural manifold

We prepared two videos. One, denoted as Supplementary Movie 1, serves to illustrate the stimuli, and presents a schematic diagram of the eight conditions. The other video, denoted as Supplementary Movie 2, demonstrates the dynamic changes in classification accuracy corresponding to the neural manifold. For additional information, please refer to the supplementary files.

### Supplementary Methods

In this Supplementary Methods, we present more detailed versions of the methods in the main text, including formulations, equations, algorithms, etc.

#### M.1. Arithmetic product as continuous version of exclusive OR (XOR)

There is correspondence between XOR and arithmetic product. Let us correspond the Boolean true value to -1.0 and the Boolean false value to 1.0 as shown in Supplementary Fig. 25. For instance, a Boolean variable X is true if and only if the corresponding real variable x = -1.0, and false if and only if x = 1.0. We find that the truth values of $X \oplus Y$ align well with the results of $x \cdot y$.

Unlike XOR operation, arithmetic product can deal with continuous values, i.e., $x, y \in \mathbb{R}$. Hence, arithmetic product can be regarded as a continuous version of XOR operation. With continuous variables, arithmetic product effectively renders a twisting operation shown in Fig. 4A.

**A**

| xor $\oplus$ | | |
|---|---|---|
| $x$ | $y$ | $x \oplus y$ |
| F | F | F |
| F | T | T |
| T | F | T |
| T | T | F |

**B**

| product | | |
|---|---|---|
| $x$ | $y$ | $x \cdot y$ |
| 1.0 | 1.0 | 1.0 |
| 1.0 | -1.0 | -1.0 |
| -1.0 | 1.0 | -1.0 |
| -1.0 | -1.0 | 1.0 |

**Supplementary Figure 25. The correspondence between XOR and arithmetic product**

(A) The truth table of XOR operation on two Boolean variables $X$ and $Y$. F: False, and T: True.

(B) The results of multiplying two real number $x$ and $y$. The color code indicates the correspondence between -1.0 and the true value, and that between 1.0 and the false value. The last column in panel B aligns with the last column in panel A according to this correspondence.

#### M.2. Explanation of support vector machine (SVM)

The SVM used in this study is a binary linear classifier that maximizes the margin between the cluster boundaries of two classes of data points in high-dimensional spaces. Formally, let $\boldsymbol{x}$ be a point in the $n$-dimensional neural space. With respect to a stimulus feature, a point $\boldsymbol{x}$ can be labelled by $y = \pm 1$ , corresponding to the positive and negative classes, respectively. The class separating hyperplane is represented by the equation:

$$f(\boldsymbol{x}) = \boldsymbol{x}^{\mathrm{T}}\boldsymbol{\beta} + b = 0,$$

where $\boldsymbol{\beta}$ is the unit vector representing the direction of the hyperplane, and $b$ is the bias along the direction from the origin of the space. The predicted class of $\boldsymbol{x}$ is determined by the sign of $f(\boldsymbol{x})$. The optimization process aims to find the optimal values of $\boldsymbol{\beta}$ and $b$ that maximize the margin. The optimized $\boldsymbol{\beta}$ gives us the classification direction.

Specifically, after mathematical derivations (Cristianini et al., 2000), for all labelled data points $(\boldsymbol{x_i}, y_i)$ the optimization is

$$\min_{\boldsymbol{\beta},b,\xi}\left(\frac{1}{2}\boldsymbol{\beta}^{\mathrm{T}}\boldsymbol{\beta} + c\sum_i \xi_i^2\right),$$

such that

$$y_i f(\boldsymbol{x_i}) \geq 1 - \xi_i,\ \ \xi_i \geq 0,$$

where $\boldsymbol{\xi} = [\xi_1, \xi_2, \ldots]$ represents the slack variables that allow soft separation. The value of the box constraint $c$ controls the softness of the separation.

Softness is introduced to prevent overfitting of the SVM classifier. A softer separation permits more classifying errors, while a harder separation allows fewer classification errors. Careful selection of the box constraint is necessary to ensure the classifier is accurate enough without being overfitted to noise.

**M.3. Algorithm of linear transformation for 3-D visualization**

Suppose we have $m$ mutually orthogonal main axes which are $n$-dimensional unit vectors $\boldsymbol{v_1}, \boldsymbol{v_2}, \ldots, \boldsymbol{v_m}$. The algorithm to generate linear transformation matrix $\boldsymbol{T}$, from the original neural space to the new coordinate system, is in Algorithm 1.

Lines 1–2 initialize a matrix $\boldsymbol{A}$ with information of the main axes embedded. Line 3 converts $\boldsymbol{A}$ into an orthogonal matrix $\boldsymbol{Q}$ so that it can be a transformation matrix. The first $m$ columns of $\boldsymbol{A}$ were generated by SVM classifications. Empirically, the first $m$ columns are approximately orthogonal (Flesch et al., 2002; Libedinsky, 2023). Accordingly, the first $m$ columns of $\boldsymbol{Q}$ are approximately parallel to the first $m$ columns of $\boldsymbol{A}$. Lines 4–9 ensure the first columns of $\boldsymbol{Q}$ and $\boldsymbol{A}$ have same instead of opposite directions. The output line follows the fact that the relation between a data point's description $\boldsymbol{b}$ in the new coordinate system and the data point's description $\boldsymbol{a}$ in the original coordinate system is $\boldsymbol{a} = \boldsymbol{Q}\boldsymbol{b}$. We can use $\boldsymbol{b} = \boldsymbol{Q}^{-1}\boldsymbol{a}$ to get the description in the new coordinate system. Since $\boldsymbol{Q}$ is an orthogonal matrix, we get the transformation matrix as $\boldsymbol{T} = \boldsymbol{Q}^{-1} = \boldsymbol{Q}^{\mathrm{T}}$.

**Algorithm 1**: **Linear transformation matrix**

**Input:** $\boldsymbol{v_1}, \boldsymbol{v_2}, \dots, \boldsymbol{v_m}$

Initialized a $n \times n$ identity matrix $\boldsymbol{I}$

Replace the first $m$ columns of $\boldsymbol{I}$ with $\boldsymbol{v_1}, \boldsymbol{v_2}, \dots, \boldsymbol{v_m}$ to get a new matrix $\boldsymbol{A}$.

Use QR decomposition $[\boldsymbol{Q}, \boldsymbol{R}] = \mathrm{qr}(\boldsymbol{A})$ in Matlab to get an orthogonal matrix $\boldsymbol{Q}$ and an upper triangular matrix $\boldsymbol{R}$.

**For** $i = 1 \dots \mathrm{m}$

    **If** the $i$th diagonal element of $\boldsymbol{R}$ is negative

        Change this element to positive

        Prefix a negative sign to the $i$th column of $\boldsymbol{Q}$.

    **End**

**End**

**Output:** $\boldsymbol{T} = \boldsymbol{Q}^{\mathrm{T}}$

#### M.4. Fitting the double-twist model to neural data

We used affine transformation to fit the double-twist model to the neural data. We first applied the low-pass filter (cutoff frequency: 2Hz) to reduce the noise. Then we computed the steady-state averages (300ms–500ms relative to stimulus onset) of the neural activities for the 8 stimuli to get 8 data centroids. The 8 data centroids $\boldsymbol{y_i}$ , $i = 1, \dots, 8$, in each 3-D subspace were stored as columns in a matrix $\boldsymbol{Y}$.

We found that 8 locations $\boldsymbol{x_i}$ on the ideal double-twisting model corresponded to the 8 data centroids in the same 3-D subspace. These locations were stored as columns in $\boldsymbol{X}$. We wanted to find a transformation $\boldsymbol{Y} = \boldsymbol{FX}$. The matrix $\boldsymbol{F}$ was computed using the least-square method as $\boldsymbol{F} = \boldsymbol{YX}^{\mathrm{T}}(\boldsymbol{XX}^{\mathrm{T}})^{-1}$. However, the least-square method generated nonzero residuals $\boldsymbol{E} = \boldsymbol{Y} - \boldsymbol{FX}$. We computed the mean of the column vectors of $\boldsymbol{E}$ as the residual vector $\boldsymbol{e}$. The matrix $\boldsymbol{F}$ and the vector $\boldsymbol{e}$ constituted the affine transformation. For any point $\boldsymbol{x}$ on the ideal double-twist model, we applied the transformation:

$$\widehat{\boldsymbol{y}} = \boldsymbol{Fx} + \boldsymbol{e}.$$

The goodness of fit between 8 data points $\boldsymbol{y_i}$ and 8 fitted locations $\widehat{\boldsymbol{y}}_{\boldsymbol{i}}$ was computed using $\mathrm{R}^2$ as

$$R^2 = \frac{\sum_{i=1}^{8} \|\hat{\boldsymbol{y}}_i - \bar{\boldsymbol{y}}\|^2}{\sum_{i=1}^{8} \|\boldsymbol{y}_i - \bar{\boldsymbol{y}}\|^2}.$$

**M.5. Binary classification**

Binary classification involves classifying stimulus conditions into two classes based on all possible classification rules. In this context, the number of all possible classification rules is $2^N$, where $N$ represents the number of stimulus conditions. As we had 8 stimulus conditions, there were $2^8 = 256$ classification rules in total. The neural states for the same stimulus condition were clustered, and thus neural space had 8 clusters in total. Accordingly, for each classification, the criterion for linear separability in the neural space was set to achieve an accuracy above 75% for each stimulus cluster, rather than considering the overall accuracy across all stimulus states. In the actual analysis, we only tested 127 classification rules because the other half of the classification rules are symmetric to them, and all conditions belonging to one category are inherently linearly separable.

In order not to underestimate the number of linearly separable classifications because of noisy stimulus, here we only used data from coherence level 7 and correct trials, where box constraint $c$ for SVM analysis was set to 1. In addition, each neuron pool consisting of varying number of neurons was randomly selected from the whole 93 neurons one at a time, and the SVM analysis was conducted 10 times for each neuron pool. In the case of the neuron pool that already included all 93 neurons, we conducted the SVM analysis only once.

**M.6. Dimensionality of artificial neural networks**

We calculated the dimensionality of the given artificial neural network with a specific diversity $d$. Firstly, we chose a number $n$ ($n \in \{2, \dots, 8\}$) increasingly from 2 to 8, and randomly selected $n$ conditions from the whole 8 conditions. Then, we performed SVM analysis on these $n$ conditions to obtain the actual number of linearly separable classifications and calculated the theoretical number of binary classifications as $2^n$. Finally, we calculated the ratio of these two numbers. In this way, we got one ratio for each $n$. The last $n$ whose ratio > 0.8 was selected as the dimensionality of this network. Using this procedure, we ran 100 simulations of the networks, obtaining 100 dimensionality for each diversity $d$. We calculated means and standard deviations of the dimensionality for different diversity $d$ and plot them as in Fig. 6H.

687
